# Cone-shaped HIV-1 capsids are transported through intact nuclear pores

**DOI:** 10.1101/2020.07.30.193524

**Authors:** Vojtech Zila, Erica Margiotta, Beata Turonova, Thorsten G. Müller, Christian E. Zimmerli, Simone Mattei, Matteo Allegretti, Kathleen Börner, Jona Rada, Barbara Müller, Marina Lusic, Hans-Georg Kräusslich, Martin Beck

## Abstract

Human immunodeficiency virus (HIV-1) remains a major health threat. Viral capsid uncoating and nuclear import of the viral genome are critical for productive infection. The size of the HIV-1 capsid is generally believed to exceed the diameter of the nuclear pore complex (NPC), indicating that capsid uncoating has to occur prior to nuclear import. Here, we combined correlative light and electron microscopy with subtomogram averaging to capture the structural status of reverse transcription-competent HIV-1 complexes in infected T cells. We demonstrate that the diameter of the NPC *in cellulo* is sufficient for the import of apparently intact, cone-shaped capsids. Subsequent to nuclear import, we detected disrupted and empty capsid fragments, indicating that uncoating of the replication complex occurs by breaking the capsid open, and not by disassembly into individual subunits. Our data directly visualize a key step in HIV-1 replication and enhance our mechanistic understanding of the viral life cycle.

## Introduction

Human immunodeficiency virus type 1 (HIV-1) is a lentivirus that infects non-dividing cells (Yamashita and Emerman, 2006). The primary targets of HIV-1 *in vivo* are immune cells, including CD4^+^ T lymphocytes and macrophages (Stevenson, 2003). HIV-1 enters the cells by fusion of the virion envelope with the cell membrane (Chen, 2019), which leads to release of the viral capsid into the cytosol. The mature HIV-1 capsid is a cone-shaped structure of ~120 by 60 nm with fullerene geometry. It is composed of ~1,200 copies of the viral CA (capsid) protein that assemble into a lattice of 250 CA hexamers (Briggs et al., 2003; Sundquist and Kräusslich, 2012). Five and seven CA pentamers incorporated at the narrow and broad end of the cone, respectively, close the capsid and induce the characteristic curvature (Ganser et al., 1999; Mattei et al., 2016). The capsid shell encases two copies of genomic single stranded RNA associated in a condensed ribonucleoprotein (RNP) complex with the NC (nucleocapsid) protein, the replication enzymes reverse transcriptase (RT) and integrase (IN) as well as other components (Welker et al., 2000). Following cytosolic entry, the viral replication complex undergoes reverse transcription of the RNA genome into double-stranded DNA and transport into the nucleus, where the viral genome integrates into that of the host cell (Engelman and Singh, 2018; Hu and Hughes, 2012).

Reverse transcription and integration are mediated by poorly characterized subviral complexes with unknown morphology termed reverse transcription complexes (RTC) and preintegration complexes (PIC), respectively (Engelman and Singh, 2018; Hu and Hughes, 2012). The fact that reverse transcription and integration are rare events in an infected cell, and the transient nature of these processes, precluded a detailed biochemical and structural characterization of RTC and PIC so far. Initially, the viral capsid was assumed to rapidly disassemble upon entry into the cytosol, but more recent evidence indicated that incoming capsids remain intact at least through the initial stages of reverse transcription (Novikova et al., 2019). The capsid structure has been suggested to play a crucial role during early replication, including intracellular trafficking, protection of RTC/PIC against innate immune sensing and import of the genome into the nucleus (Ambrose and Aiken, 2014; Campbell and Hope, 2015; Hilditch and Towers, 2014; Yamashita and Engelman, 2017). The transport of the HIV-1 RTC/PIC towards the nucleus relies on microtubules (MTs) (Arhel et al., 2006; Malikov et al., 2015; McDonald et al., 2002) and requires the association of CA with dynein and kinesin-1 motors mediated by MT-associated adaptor proteins (Dharan et al., 2017; Fernandez et al., 2015; Malikov et al., 2015). Perinuclear movements and docking of subviral complexes to the nuclear envelope might be mediated by the actin cytoskeleton (Arhel et al., 2006), but also the relocation of NUP358/RanBP2 from the NPC to cytosolic CA mediated by kinesin-1 was observed to precede nuclear import of HIV-1 PIC (Dharan et al., 2016).

Nuclear import of the PIC and integration of HIV-1 genomic DNA into the host genome are interconnected processes. They are essential for productive HIV-1 infection, but also for the establishment of the HIV-1 latent reservoir, a silenced pool of replication-competent proviruses persisting in resting CD4^+^ T cells and resistant to antiretroviral therapy (ART) (Lusic and Siliciano, 2017). Active nuclear import of the PIC is facilitated by a nuclear localization signal (NLS) in the cyclophilin A (CypA)-binding loop of CA that is recognized by the nuclear transport receptor transportin (Fernandez et al., 2019). Several nucleoporins (NUPs), most notably the FG-repeat containing NUP358 and NUP153 were also reported to facilitate nuclear entry of HIV-1 (Brass et al., 2008; Di Nunzio et al., 2012; König et al., 2008). While NUP358 may mediate docking of the complex to the cytoplasmic face of the NPC (Dharan et al., 2016), it remains unclear how HIV-1 complexes cross the central channel. Once viral complexes have reached the nuclear basket, they can interact with NUP153 *via* a hydrophobic pocket on CA hexamers to promote the final steps of PIC translocation (Campbell and Hope, 2015; Matreyek et al., 2013; Price et al., 2014). The nuclear protein Cleavage and Polyadenylation Specificity Factor 6 (CPSF6) was suggested to compete with NUP153 for the common binding site on CA (Matreyek et al., 2013), resulting in the release of the PIC into the nucleus (Bejarano et al., 2019). Importantly however, the HIV-1 capsid is about ~60 nm wide at the broad end of the cone (Briggs et al., 2003; Mattei et al., 2016), which considerably exceeds the inner diameter of NPCs as seen in cryo-EM structures obtained from isolated nuclear envelopes, which is only ~40 nm (von Appen et al., 2015). While this discrepancy in size suggests that the PIC cannot pass the central channel without breaking the CA lattice, recent structural analyses in intact cells indicated that NPCs may occur in a dilated conformation under certain circumstances (Beck and Baumeister, 2016). However, the NPC structure has never been studied in the relevant cell types under conditions of HIV-1 infection; thus, the physiological relevance of the latter observation for HIV-1 nuclear import remains unknown.

At least a partial dissociation of the HIV-1 capsid lattice (uncoating) is a prerequisite for the release of the PIC prior to genome integration, but the timing, cellular location and extent of capsid uncoating are still not clear and might be cell type specific. Several models have been put forward, including i.e. gradual uncoating with concomitant reverse transcription during cytosolic trafficking; reverse transcription within largely intact capsids followed by their uncoating at the NPC; or several, spatially separated uncoating steps that are finalized only in the nucleus (Campbell and Hope, 2015; Novikova et al., 2019; Zhou et al., 2011). Recent data indicate that HIV-1 nuclear import precedes the completion of reverse transcription (Dharan et al., 2020). Tracking experiments of individual HIV-1 complexes by fluorescent microscopy in living cells supported capsid uncoating within the cytosol (Mamede et al., 2017), at the NPC (Francis and Melikyan, 2018) or inside the nucleus (Burdick et al., 2020), and variable amounts of CA have been detected on nuclear HIV-1 PICs in different cell types (Bejarano et al., 2018, 2019; Chen et al., 2016; Chin et al., 2015; Hulme et al., 2015; Peng et al., 2014; Stultz et al., 2017; Zhou et al., 2011). However, whether the lattice structure remains intact in these complexes or alternatively, CA remains associated with the RTC/PIC despite lattice disassembly, remains unknown.

To address these questions, we employed 3D correlative fluorescence light and electron microscopy (CLEM), cryo-electron tomography (cryo-ET) and subtomogram averaging to examine the ultrastructure of early HIV-1 replication complexes during cytosolic transport and nuclear import in an infected human CD4^+^ T cell line. We found that subviral HIV-1 complexes maintained their typical cone-shape in the cytosol upon association with microtubules, and when docked to NPCs. The overall shape and hexagonal lattice of the capsid remained detectable even during transport through NPCs, which is possible because the NPC scaffold was dilated with respect to previous structural analysis of isolated nuclear envelopes. HIV-1 complexes inside the nucleus, on the other hand, had lost their cone shape and lacked interior dense material. Our findings suggest that intact HIV-1 capsids pass the nuclear pore, and disruption of the capsid lattice occurs inside the nucleus and facilitates the release of the PIC.

## Results

### An experimental system for the ultrastructural analysis of HIV-1 post-entry complexes

Chemical fixation is commonly used to inactivate infectious particles. However, it negatively affects the structural preservation of the sample. In order to study the ultrastructure of early HIV-1 replication complexes in cryo-immobilized infected cells at biosafety level 1, we constructed an HIV-1 expression plasmid (pNNHIV) for production of the non-infectious RT-competent HIV-1 derivative NNHIV (Figure 1A; Figure S1A, B). Two point mutations were introduced into the IN active site to prevent integration of the proviral genome. Furthermore, the viral accessory protein Tat was truncated in order to block transactivation of HIV-1 transcription. Digital droplet PCR (ddPCR) confirmed that NNHIV reverse transcription kinetics in infected SupT1-R5 cells (Figure 1B, C) was similar to that previously reported for wild-type HIV-1 (Zila et al., 2019). Late RT products of NNHIV were detected from 3 h postinfection (p.i.) onwards and peaked at 12 h p.i. with the majority of late RT products synthesized between 3 and 6 h p.i. (Figure 1B), which is consistent with our previous findings (Zila et al., 2019). NNHIV 2-LTR (long terminal repeat) circles, a surrogate for replication complexes transported into the nucleus, were detected from 6 h p.i. onwards; the strong accumulation of 2-LTR circles in NNHIV infected cells compared to those infected with the wild-type virus (Figure 1C) was indicative of the block in NNHIV genome integration. No specific ddPCR products were detected upon NNHIV infection in the presence of the nucleoside RT inhibitor Efavirenz (EFV) (Figure 1B, C). We conclude that NNHIV undergoes reverse transcription and nuclear import with dynamics similar to wild-type HIV-1.

**Figure 1.**
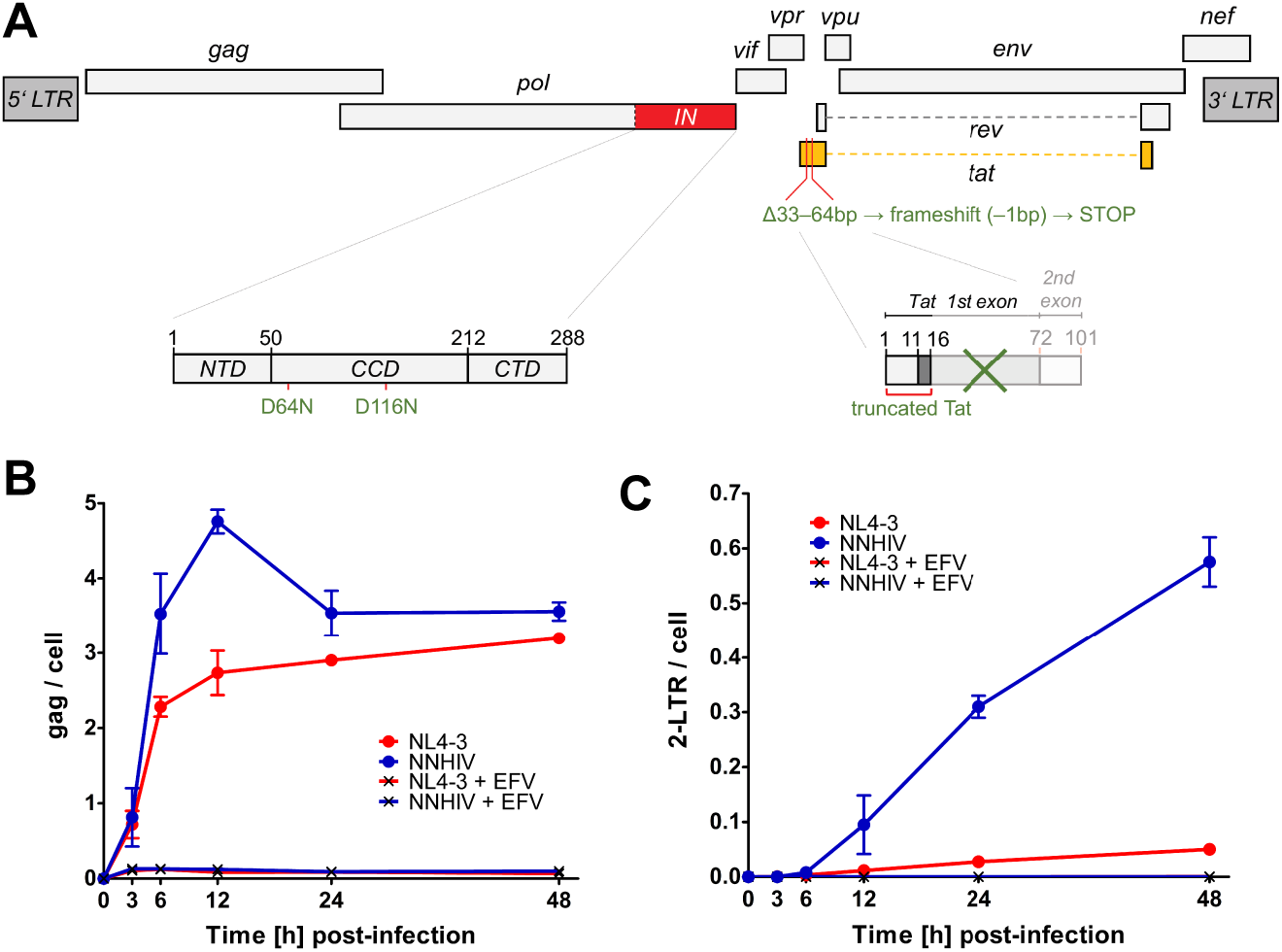
Generation of reverse transcription-competent, non-infectious HIV-1 derivative for EM studies. **(A)** The genome of NNHIV, a reverse transcription competent, IN- and Tat-defective HIV-1 derivative. The integrase coding sequence (IN; red) and *tat* gene (orange) are highlighted. The position of two mutated amino acid residues (D64N and D116N) introduced in IN catalytic core domain (CCD), the truncated Tat protein as a result of the frame shift (−1 bp) introducing a premature stop codon in *tat* are enlarged. NTD, N-terminal IN domain; CTD, C-terminal IN domain. **(B, C)** Quantification of NNHIV RT products using digital droplet PCR (ddPCR). SupT1-R5 cells were infected with equal amounts of NNHIV or wild-type HIV-1_NL4-3_ virions (NL4-3) in the absence or presence of EFV. Cells were harvested at the indicated time points and HIV-1 RT products were quantitated in cell lysates by ddPCR. Copy numbers of late RT products (B) and 2-LTR circles (C) were normalized to the copy numbers of the housekeeping gene RPP30. Graphs show mean values and SD from triplicate samples. Data for NL4-3 from (Zila et al., 2019), derived from the same experiment, are shown for comparison.

We recently established a fluoresce microscopy approach to discriminate post-fusion HIV-1 complexes in the cytosol from intact virions either at the plasma membrane or inside of endosomes (Zila et al., 2019). SupT1-R5 cells were infected with HIV-1 particles carrying fluorescently labeled IN (Albanese et al., 2008) as a marker for the HIV-1 core and stained with the fluorescently tagged endocytic probe mCLING (Revelo et al., 2014) to label the plasma membrane and endosomes. IN positive objects within the cell that lacked the mCLING membrane marker were defined as cytosolic HIV-1 post-fusion complexes. To utilize this approach for electron microscopy (EM) studies, optimal preservation of both fluorescent signals throughout the sample preparation for ultrastructural analysis was obtained by using a combination of mCLING.Atto647N and IN fused to mScarlet. Control experiments confirmed that incorporation of exogenously expressed IN.mScarlet did not have a major effect on viral infectivity (Figure S1B–D), similar to what had been observed for IN.eGFP under similar conditions (Peng et al., 2014).

We next established a workflow for the identification of HIV-1 post-entry complexes using fiducial-based on-section CLEM (Kukulski et al., 2011) in combination with electron tomography (ET) (Figure S2). NNHIV particles carrying IN.mScarlet were adhered to SupT1-R5 cells for 90 min at 16°C. The low temperature prevents both HIV-1 membrane fusion (Henderson and Hope, 2006; Melikyan et al., 2000) and endocytosis (Punnonen et al., 1998; Weigel and Oka, 1981). The plasma membrane was stained with mCLING.Atto647N for an additional 10 min at 16°C and samples were then shifted to 37°C to initiate virus entry (Zila et al., 2019) (Figure 2A). To maximize cytosolic entry events, we incubated cells at 37°C for 90 min (Zila et al., 2019). Infected cells were then subjected to high pressure freezing (HPF) and freeze substitution (Kukulski et al., 2011). Spinning disc confocal microscopy (SDCM) of 250 nm-thick resin sections revealed bright fluorescence of both probes at the plasma membrane and in endosomes. As expected, we detected mCLING-negative IN.mScarlet foci in the cell interior, which were indicative of post-fusion complexes (Figure S2B). Such foci identified in EM sections of cells were selected as regions of interest (ROI) for correlative ET imaging. Tomograms obtained from ROIs revealed the presence of dense conical structures within the cytosol, visually distinct over the dense cellular background (Figure S2B–D).

**Figure 2.**
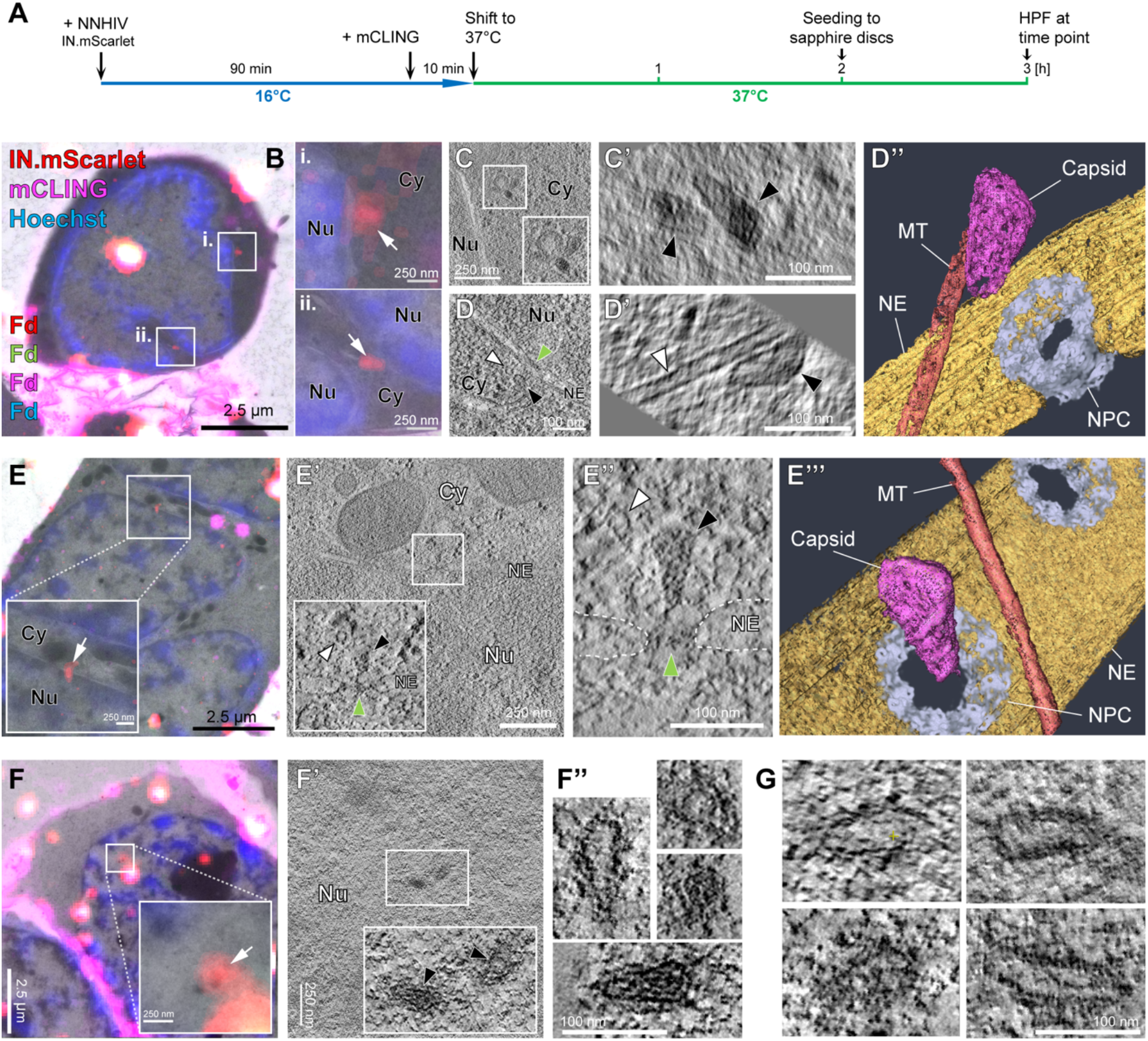
HIV-1 capsids are intact in the cytosol and adjacent to the NPC but morphologically altered inside the nucleus of T cells. SupT1-R5 cells were infected with IN.mScarlet carrying NNHIV for 90 min at 16°C. To identify post-fusion subviral complexes, cells were stained with mCLING.Atto647N for additional 10 min at 16°C. Cells were subsequently incubated in the presence of mCLING.Atto647N for 3 h at 37°C prior to cryoimmobilization by high pressure freezing (HPF) and freeze substitution. **(A)** Schematic illustration of the experiment. **(B–D)** Morphology of HIV-1 complexes in the cytosol. **(B)** CLEM overlay of a 250-nm thick resin section of the cell stained with mCLING.Atto647N (far-red; magenta), post-stained with Hoechst (blue) and decorated with multi-fluorescent fiducials (Fd) for correlation. Enlarged regions indicate presence of mCLING-negative IN.mScarlet signals (red) in two positions (i. and ii.) within the cytosol (white arrows). **(C, D)** Slices through a tomographic reconstruction at the correlated positions i. (C) and ii. (D), with enlarged views highlighting cone-shaped capsids (black arrowheads) in the cytosol (C’) and in proximity to MTs (D’; white arrowhead). **(D”)** Same as in (D’) but displayed segmented and isosurface rendered. MT red; capsid, magenta; NE, yellow; NPC, cyan (cryo-EM map of NPC: this study). **(E)** CLEM overlay enlarged in the inset shows mCLING-negative signal of IN.mScarlet (red) located at the nuclear envelope (white arrow). **(E’, E’’)** Correlated position within an electron tomogram with enlarged inset (E’) and slice through a tomographic reconstruction (E’’) showing a MT-associated capsid docking to the NPC. Black, white and green arrowheads indicate the capsid, microtubule cross section and NPC, respectively. Dashed lines outline the nuclear membrane. **(E’’’)** Same as in (E’’) but segmented and isosurface rendered. Color code as in (D’’). **(F-F’’)** Same as in (E-E”) but for NNHIV complexes captured inside of the nucleus, highlighting the morphology of four clustered, capsid-related structures. **(G)** Further representative examples of nuclear NNHIV complexes captured in different cells. Cy, cytosol; Nu, nucleus; NE, nuclear envelope; NPC, nuclear pore complex.

### Cone-shaped HIV-1 capsids dock to the NPC

To examine NNHIV post-entry and NPC docking events (Zila et al., 2019), SupT1-R5 cells infected and stained as described above were high pressure frozen at 3 h post temperature shift (Figure 2A). We acquired tomograms in a total of 45 positions of correlated ROIs from two independent experiments. From this data set we identified 26 structures that resembled intact HIV-1 capsids in the cytosol or adjacent to nuclear pores. Their morphology closely matched that of mature capsids within HIV-1 virions, including an accumulation of dense material within the shell indicating the presence of condensed RNP or reverse transcription intermediates (Figure 2B, C). The majority of structures (22/26; 85%) was cone-shaped with an average length of 112 ± 12 and average width of 53 ± 7, similar to the dimensions determined for mature HIV-1 capsids by cryo-ET (Briggs et al., 2003). Most structures (21/26; ~80%) were found associated with microtubules, including those in the proximity of NPCs (n = 7; average distance to the NPC inner ring 33 ± 9 nm; Figure 2D–D”’; Video S1). Two of the docking capsids were oriented perpendicular to the NPC with their narrow end pointing towards the central channel (Figure 2E–E”’; Video S2). Together these data indicate that apparently intact HIV-1 capsids associated to microtubules dock to the NPC in infected T cells.

### HIV-1 capsid morphology is altered in the nucleoplasm

In addition to cytosolic structures described under above conditions, we also detected labeled complexes inside of nuclei. A total of 11 individual structures, as well as 4 structures in close proximity to each other were identified by ET in sections of eight different cells. Their morphology clearly differed from the structures observed in the cytosol (Figure 2F, G; Video S3). With one exception (Figure 2F”), the nuclear structures appeared to be open and their interior was devoid of dense material, suggestive of separation of the nucleoprotein complex from the broken capsid shell. The majority of empty shells had a tubular shape, but cone-like remnants were also observed (Figure 2F, G; Video S3).

### Cone-shaped HIV-1 capsids can enter the central channel of the NPC

The data described above revealed that HIV-1 capsids underwent a structural change on the way from the cytoplasmic NPC docking site to the nucleoplasm, but did not allow to pinpoint the exact stage or site where this change occurred. In order to characterize the ultrastructure of HIV-1 complexes during nuclear import, we performed CLEM and cryo-ET analyses under conditions that enrich for viral complexes at nuclear pores. For this, we employed an NNHIV-derivative carrying a mutation in CA (A77V) previously reported to prevent the interaction with CPSF6 (Saito et al., 2016). Impairing CA-CPSF6 interaction results in steady-state accumulation of PICs at nuclear pores without a major effect on virus infectivity in monocyte derived macrophages or SupT1-R5 T cells (Bejarano et al., 2019; Zila et al., 2019). SupT1-R5 cells were incubated with A77V NNHIV particles carrying IN.mScarlet. After low temperature adsorption and mCLING staining (Figure 2A), cells were incubated for 15 h at 37°C to allow for accumulation of viral complexes at NPCs, prior to high pressure freezing. Tomograms of intracellular ROIs from 7 different cells sections revealed 14 conical capsids in the cytosol or associated with nuclear pores. Strikingly, several cone-shaped A77V capsids were visualized deep inside the NPC channels, exposing their narrow ends to the nucleoplasm (n = 3; Figure 3A–A”, Video S4). These capsids contained dense material inside presumed to correspond to the viral nucleoprotein complex (Figure 3A”, Figure S3B).

**Figure 3.**
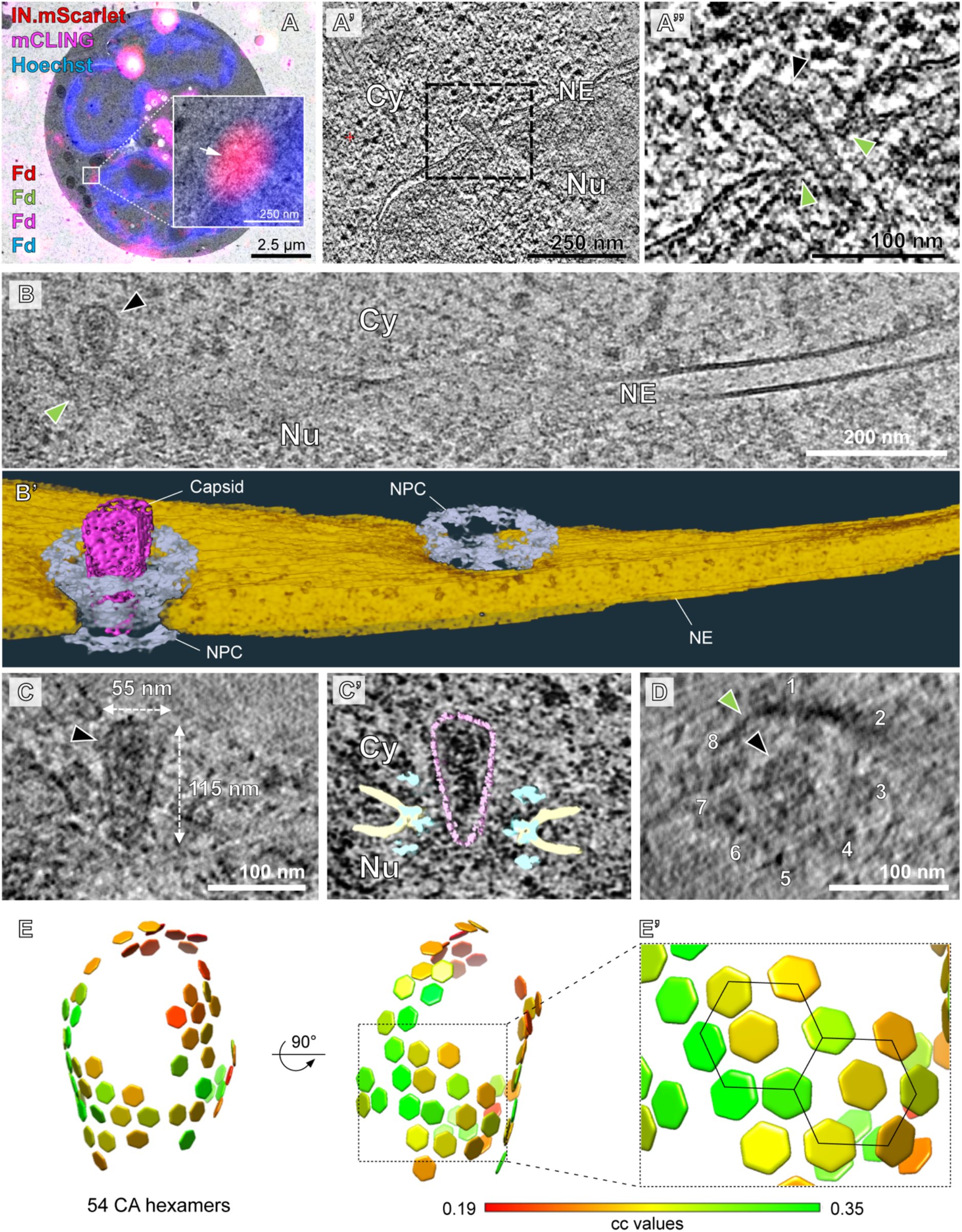
Intact CA-A77V capsids can deeply penetrate into the central channel of the NPC. SupT1-R5 cells were infected with IN.mScarlet carrying NNHIV-A77V CA mutant for 15 h at 37°C prior to high pressure freezing (for CLEM) or plunge freezing (for cryo-ET). **(A-A’’)** Plastic section CLEM of a cell stained with mCLING.Atto647N (magenta), post-stained with Hoechst (blue) and decorated with multi-fluorescent fiducials (Fd) for correlation. **(A)** Overlay of a fluorescence image with the correlated electron micrograph. The enlarged region displays the position of the IN.mScarlet signal (red) at the nuclear envelope (white arrow). **(A’)** Slice through a tomographic reconstruction at the correlated position. The boxed region is shown enlarged in (A’’) highlighting an apparently intact capsid (black arrowhead) deep inside the central channel of the NPC (green arrowheads), exposing its narrow end towards the nucleoplasm. Cy, cytosol; Nu, nucleus; NE, nuclear envelope. **(B–D)** Cryo-electron tomography of cryo-FIB milled SupT1-R5 cells infected with NNHIV-A77V. **(B)** Slice through a tomographic reconstruction showing a capsid (black arrowhead) localized inside of the NPC (green arrowhead; black box). **(B’)** Same as in (B) but displayed segmented and isosurface rendered. Capsid, magenta; NE, yellow; NPC, cyan (cryo-EM map of NPC: this study). **(C)** Enlarged and rotated view of the HIV-1 capsid (black arrowhead) framed in (B), dimensions are indicated. **(C’)** Same as (C) but superimposed with structural models of capsid (magenta; cryo-EM density map of hexameric unit: EMD-3465 (Mattei et al., 2016)) and the NPC (cyan; cryo-EM map of NPC: this study). NE in yellow. **(D)** Same as (C), but displayed in top view. The broad end of the capsid cone and fused nuclear membranes are indicated by the black and green arrowhead, respectively. The eight-fold symmetry of NPC is highlighted with numbers. **(E)** Distribution of CA hexamers along the surface of the same capsid as shown in (B-C) as identified by subtomogram averaging, cross-correlation (cc) values are shown color-coded. 54 CA hexamers were identified and visualized using the Place Object plug-in (Qu et al., 2018) for USCF Chimera (Pettersen et al., 2004). The hexagonal lattice is clearly apparent. (**E’**) Black lines highlight the regular arrangement of six CA hexamers surrounding a seventh, central CA hexamer.

To investigate viral complexes in the process of nuclear entry at the best possible structural preservation, we used focused-ion beam (FIB) milling to prepare thin cryo-lamellae of infected cells. Since cryo light microscopy of these lamellae turned out to be very challenging, we chose a brute force approach and acquired ~250 tomograms of nuclear envelopes observed in cryolamellae. The resulting reconstructions contained ~100 NPCs, and four structures that resembled the viral complexes observed in the CLEM-ET data sets were detected in close proximity of or within an NPC (Figures 3B–D; Figure S3; Video S5). The structures displayed the typical conical shape and size of ~115 nm in length and ~55 nm in width. Inside, they contained highly dense material. The cone-shaped capsid entered into and penetrated with its narrow end beyond the central channel of the NPC (Figures 3B’, C’). The tip of the cone reached to the level of the nuclear ring, the region where NUP153 resides (Figure 3C’). In tomographic slices at the level of the NPC central channel (Figure 3D), the individual spokes of the inner ring were resolved and comfortably accommodated the capsid in between them.

To address whether the capsid-like structures in the cytosol and at the NPC contained a hexagonal lattice comparable to that of mature HIV-1 cores in intact virions we used subtomogram averaging, as previously described (Mattei et al., 2016). During iterative averaging the subtomograms converged into regular hexameric lattices in which six adjacent CA hexamers surround one central CA hexamer in a regular fashion (Figure 3E, E’; Figure S4A–D). In comparison to the previous data obtained from isolated virions (Mattei et al., 2016), lattice information was recovered for less of the capsid surface. This finding might be interpreted as a partial perturbation of the hexagonal lattice. However, the clearly defined capsid edge and the well-preserved overall cone shape visible in the tomograms suggest that rather technical parameters, such as the reduced signal to noise ratio due to specimen thickness of the FIB-lamellae and the crowded cellular environment, have resulted in an incomplete lattice recovery during subtomogram averaging. We conclude that cone-shaped HIV-1 capsids containing the genomic material, with an either completely or largely intact lattice, can enter the central channel of the NPC.

### HIV-1 capsids are disrupted upon nuclear entry

We next examined subviral complexes that had passed the central channel of the NPC. Previous studies revealed that in the absence of CPSF6 binding, virus infectivity is retained in nondividing cells (Achuthan et al., 2018; Bejarano et al., 2019; Saito et al., 2016). Upon depletion of CPSF6, PICs are targeted to transcriptionally repressed, lamina-associated heterochromatin (Achuthan et al., 2018). At the same time, the perturbation of CPSF6 binding to the CA hexamer by either the A77V mutation or CPSF6 depletion, resulted in partial co-localization of CA with the basket nucleoporin NUP153 (Bejarano et al., 2019), suggesting that viral complexes may reach the nucleoplasm and be retained close to the nuclear basket. Here, we identified A77V HIV-1 complexes in close proximity of NPCs by both, plastic-embedding CLEM-ET and cryo-ET analysis (average distance from closest contour to the NPC inner ring was 26 nm ± 10; n = 9) (Figure 4; Figure S3 and Video S6), suggesting that they are still engaged in interactions with the nuclear pore.

**Figure 4.**
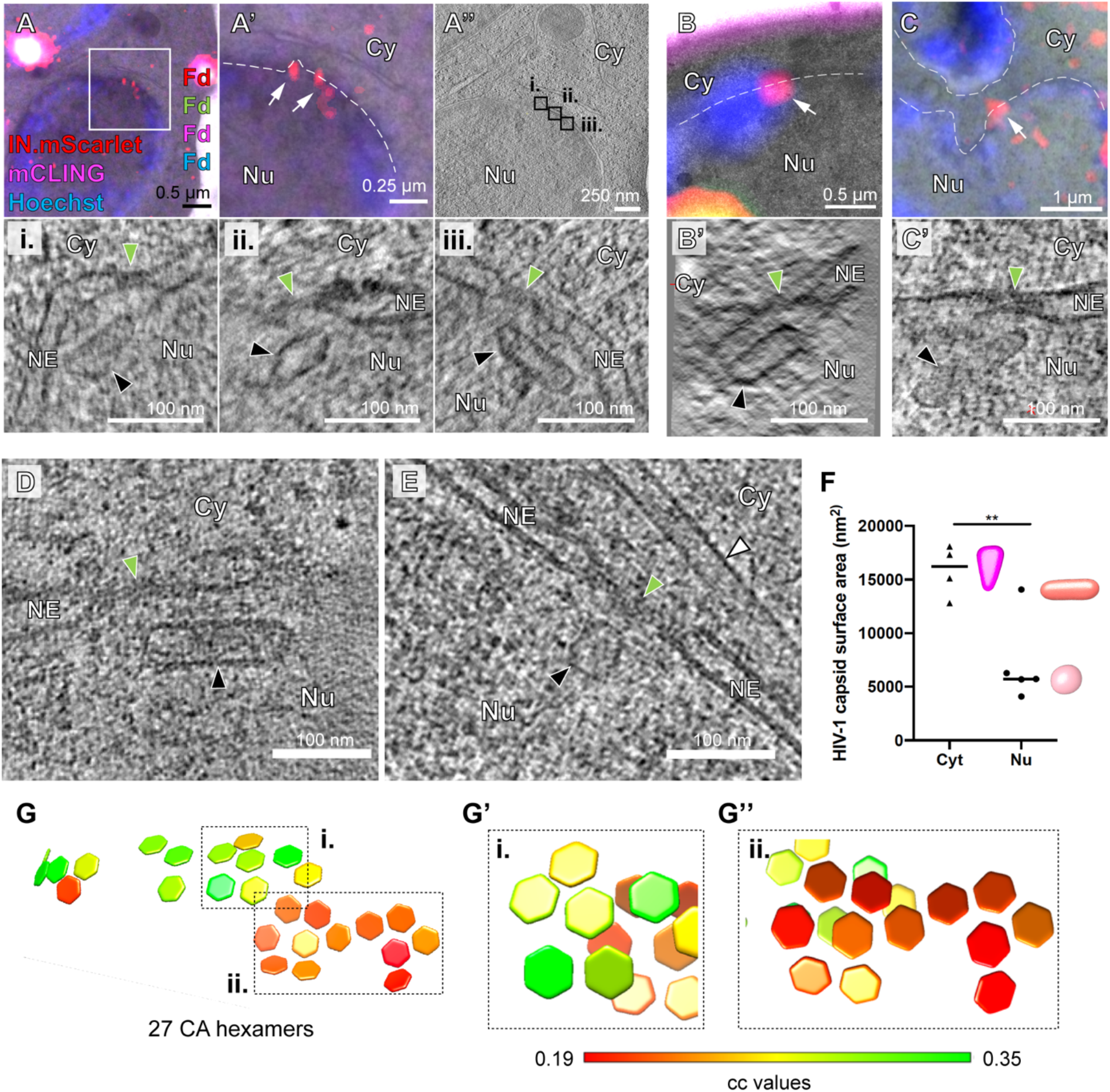
CA-A77V structures accumulated at the nuclear basket are morphologically altered. SupT1-R5 cells were infected with IN.mScarlet carrying NNHIV-A77V CA for 15 h at 37°C, prior to high pressure freezing (for CLEM) or plunge freezing (for cryo-ET). **(A-C)** Capsid-related structures at the nuclear basket region of the NPC visualized by CLEM. Dashed lines outline the nuclear envelope. **(A, A’)** CLEM overlay (A) with enlargement (A’) showing the position of IN.mScarlet signal (red; white arrow) at the nuclear envelope (empty white arrowheads) in an EM section stained with mCLING.Atto647N (magenta), post-stained with Hoechst (blue) and decorated with multi-fluorescent fiducials (Fd) for correlation. **(A’’)** Slice through a tomographic reconstruction at the correlated position shown in (A, A’). The features i.-iii. that are shown enlarged in the bottom panel are framed in black and contain three different capsid-related structures at the nuclear basket region. **(B-C’)** Same as (A-A’’) showing two further capsid-related structures from two different cells. Capsids appear tube-shaped and empty. **(D-G’’)** Cryo-ET and subtomogram averaging of NNHIV-A77V in infected SupT1-R5 cells. **(D-E)** Slices through tomographic reconstructions showing capsid-related structures localized at the nuclear basket region of the NPC. Black and green arrowheads in panels A-E indicate nuclear capsid-related structures and NPCs, respectively. **(F)** Surface area of the HIV-1 capsids identified in the cytosol (Cyt) (n = 4) and inside the nucleus (Nu) (n = 5) in the cryoelectron tomograms. Each capsid was manually segmented in IMOD and the respective surface area was computed using MATLAB. Statistical significance was assessed by unpaired twotailed Student’s t test; **p=0.0064. Representative models for three shape-based classes of nuclear structures, such as conical (magenta), tubular (salmon) and spherical (pink) are shown in the graph. **(G)** Same as in Figure 3E but for the capsid-like structure shown in (D). 27 CA hexamers were identified along the surface by subtomogram averaging. **(G’)** CA pentamer and **(G’’)** two incomplete consecutive CA hexamers.

The respective structures identified by CLEM in tomograms of plastic sections appeared morphologically altered (Figure 4A–C) as observed for wild-type complexes inside the nucleus. Most of the visualized structures had lost their cone shape, appeared partially open and were devoid of dense material presumed to correspond to the viral genome (Figure 4, compare to Figure 3). We next examined our large scale cryo-ET data set for nuclear structures. Segmentation and quantification of the tomograms identified five tube-shaped structures that visually resembled the CLEM data; four of these had only ~1/3 of the surface area compared to the conical capsids (Figure 4F). Subtomogram averaging of these four rather small particles identified only few positions with high cross-correlation with CA hexamer that did not converge into an overall hexagonal lattice. The remaining particle was also tube-shaped but considerably larger and contained some dense material inside (Figure 4D). Subtomogram averaging of its tubular core identified lattice elements (Figure 4G’-G”; Figure S4E). However, only 27 CA hexamers on the surface properly converged, much less than what was detected on the cytoplasmic structures (Figure 4, compared to Figure 3), possibly suggesting higher disorder of the lattice architecture. Taken together, the data suggest that capsid disassembly should not be conceived as immediate dissolution of the lattice into individual subunits after nuclear entry, but rather as partial disruption of the capsid that allows for the release and dissociation of the viral genome from capsid remnants, possibly due to mechanical strain. These findings support a model in which disruption of the capsid lattice occurs subsequent to translocation through or upon departure from the central channel of the NPC.

To confirm our observations independently of the A77V mutation, we infected SupT1-R5 cells with NNHIV under conditions of CPSF6 silencing. For this, we transduced cells with adeno-associated virus (AAV) vectors expressing a combination of three shRNAs targeting CPSF6, or a non-silencing shRNA. CPSF6 immunofluorescence intensities quantitated by flow cytometry revealed efficient downregulation of CPSF6 (~95%), while cell viability was not impaired (Figure S5A–C). The observed intracellular localization, the efficiency of nuclear import and the infectivity of HIV-1 upon CPSF6 knock-down in SupT1-R5 cells (Figure S5D–F) were comparable to that of the A77V mutant without knock-down (Zila et al., 2019). For CLEM, AAV-transduced cells were infected with wild-type CA-carrying NNHIV (labeled with IN.mScarlet) for 15 h prior to high pressure freezing. ET in proximity to the nuclear envelope revealed capsids docking to NPCs. Similar to the results obtained for the A77V mutant, cone-shaped capsids penetrating the NPC channel and empty, capsid-like structures at the nuclear basket were observed (Figure 5A–C, Video S7). These data reinforce the notion that HIV-1 capsids do not disassemble into individual subunits, but rather are disrupted after passage through the NPC releasing the PIC from a morphologically altered residual capsid structure.

**Figure 5.**
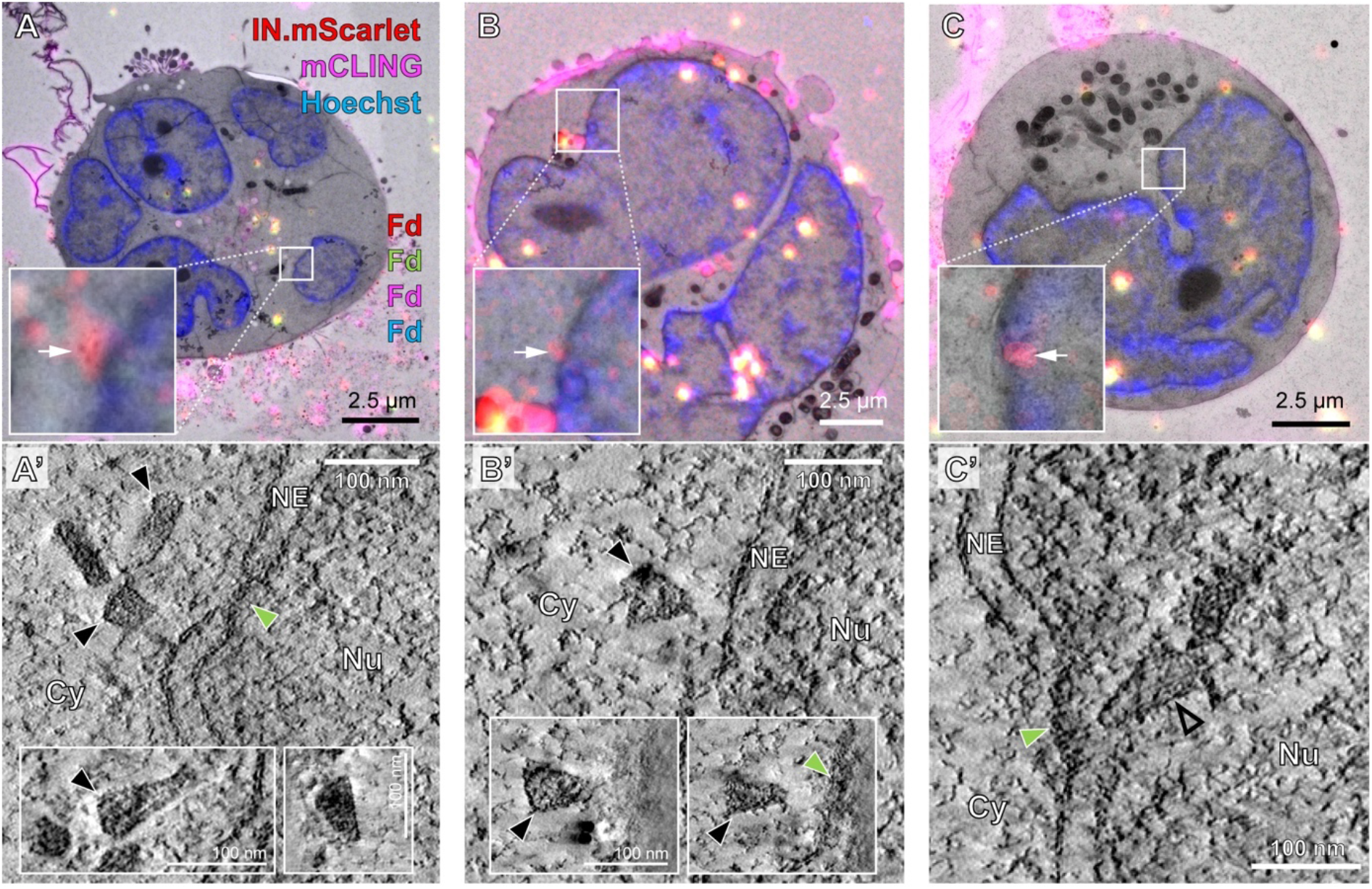
Electron tomography of NPC associated HIV-1 complexes upon CPSF6 knockdown. AAV-transduced SupT1-R5 cells were infected with IN.mScarlet-carrying NNHIV (wild-type CA) particles for 15 h at 37°C, prior to high pressure freezing. **(A-C)** CLEM overlays with enlarged regions indicate the positions of IN.mScarlet signal (red; white arrows) at the nuclear envelope in resin sections of cells stained with mCLING.Atto647N (magenta), post-stained with Hoechst (blue) and decorated with multi-fluorescent fiducials (Fd) for correlation. **(A’)** Slices through tomographic reconstructions of three different HIV-1 capsids proximate to the same NPC. Different orientations of the capsid structures are shown in the insets. Capsids at the cytoplasmic face of the NPC display a conical shape and a dense interior. **(B’)** Same as (A’) but showing an NNHIV capsid docking to the NPC. **(C’)** Same as (A’) but showing an empty capsid-related structure (empty black arrowhead) at the nucleoplasmic side of the NPC. Cy, cytosol; Nu, nucleus; NE, nuclear envelope.

### The NPC diameter is sufficient for nuclear import of intact capsids

The central channel diameter of the NPC was reported to be ~40 nm (von Appen et al., 2015), which is sufficiently wide to allow transport of basically any large cellular cargo, but too narrow to allow passage of intact HIV-1 capsids. However, the data described above revealed that largely intact HIV-1 capsids can penetrate into and pass through the NPC central channel. This observation prompted us to revisit the architecture of the NPC *in cellulo* under conditions relevant to infection. Previous structural analyses (von Appen et al., 2015) were performed using nuclear envelopes purified from human cells in which mechanical tension is relieved due to sample preparation. In order to analyze the architecture of the NPC in HIV-1 infected T cells *in situ*, we extracted 99 NPCs and 792 asymmetric units from our cryo-electron tomograms (Figure 6A) and subjected them to subtomogram averaging. The resulting cryo-EM map with a resolution of ~37 Å captures the native conformation of actively transporting NPCs in HIV-1 infected T cells (Figure S6). The overall NPC architecture appeared to be organized as previously described (Beck and Hurt, 2017) (Figure 6B, C). However, in line with other studies conducted in intact cells (Beck and Baumeister, 2016; Mahamid et al., 2016), the NPCs appeared dilated in comparison to isolated nuclear envelopes and displayed an average diameter of ~64 nm (Figure 6C, D). These findings indicate that the NPC structure observed under the relevant conditions, namely in infected T cells *in situ*, is representative of the transporting state (Beck and Baumeister 2016), while the constricted state observed in isolated nuclear envelopes (von Appen et al, 2015) may be more relevant to stress conditions (Zimmerli, Allegretti et al., attached for review process). In conclusion, our data show that the inner diameter of the central channel exceeds the dimensions of the broad end of intact HIV-1 capsids (~55–60 nm) by ~4–9 nm, rendering the nuclear entry of intact capsids geometrically possible.

**Figure 6.**
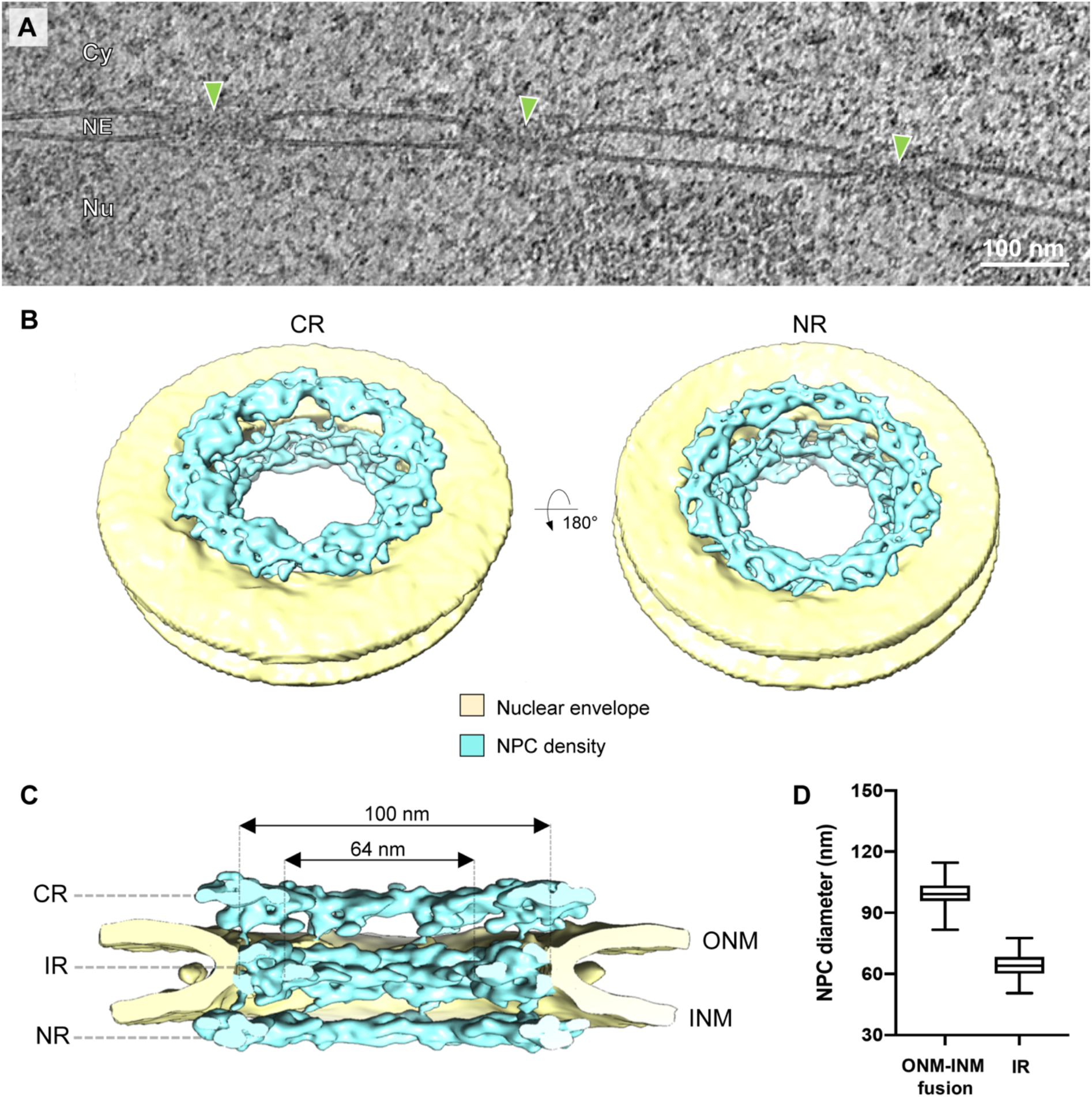
The NPC scaffold is dilated in HIV-1 infected SupT1-R5 cells. **(A)** Slice through a representative cryo-electron tomogram as used for structural analysis, containing a nuclear envelope with three NPCs (green arrowheads). Cy, cytosol; Nu, nucleus; NE, nuclear envelope. **(B, C)** Overall *in cellulo* architecture of the NPC in SupT1-R5 cells. **(B)** Isosurface rendering of the cryo-EM map of the NPC (cyan) in infected SupT1-R5 cells at ~37 Å resolution. The cytoplasmic ring (CR, left) and nuclear ring (NR, right) are visible, nuclear membrane in yellow. **(C)** Same as (B) but shown as cut-open view. CR, NR, inner ring (IR), outer nuclear membrane (ONM) and inner nuclear membrane (INM) are labelled. **(D)** Box plot showing the distribution of diameters of individual NPCs measured either at the membrane fusion site or at the relevant opening of central channel at the inner ring (IR) (n = 90). The median values were 100 nm and 64 nm, respectively.

## Discussion

Here, we have visualized the ultrastructure of HIV-1 post-entry complexes within the native cellular environment of the infected T-cell line SupT1-R5 by 3D CLEM, cryo-ET and subtomogram averaging. The combination of these methods allowed us to characterize the architecture of the capsid, including the hexameric CA lattice *in situ*. Using cryo-ET and subtomogram averaging, we further examined the native conformation of NPC in HIV-1 infected SupT1-R5 cells. Taken together, our data point to the scenario schematically outlined in Figure 7. Following cytosolic entry, intact cone-shaped HIV-1 capsids travel along microtubules towards the nuclear periphery. The subsequent nuclear import of capsids is three-staged. (i) Intact capsids dock to NPCs with the pentamer-rich ends of the capsid, preferably the narrow end. Here they encounter the FG-repeats and Cyp domain of NUP358 bound to the cytoplasmic face of NPCs. (ii) Subsequently, intact capsids penetrate deeply into the central channel of the NPC where they are exposed to a very high local concentration of FG-NUPs of the NUP62 complex. Although this environment is spatially confined, the diameter of the NPC central channel as determined *in cellulo* is physically compatible with translocation of the intact HIV-1 capsid. Up to this stage, the hexagonal lattice and the typical shape of HIV-1 capsids are clearly detectable. (iii) Upon departure from the NPC central channel, capsids encounter NUP153 and CPSF6. At this stage, which can still be conceived as part of the actual nuclear import process, disrupted capsids are detected. The cone shape is lost in many particles and smaller capsid fragments are observed. These structures lack the interior dense material, i.e. the PIC has been released for integration into the host genome.

**Figure 7.**
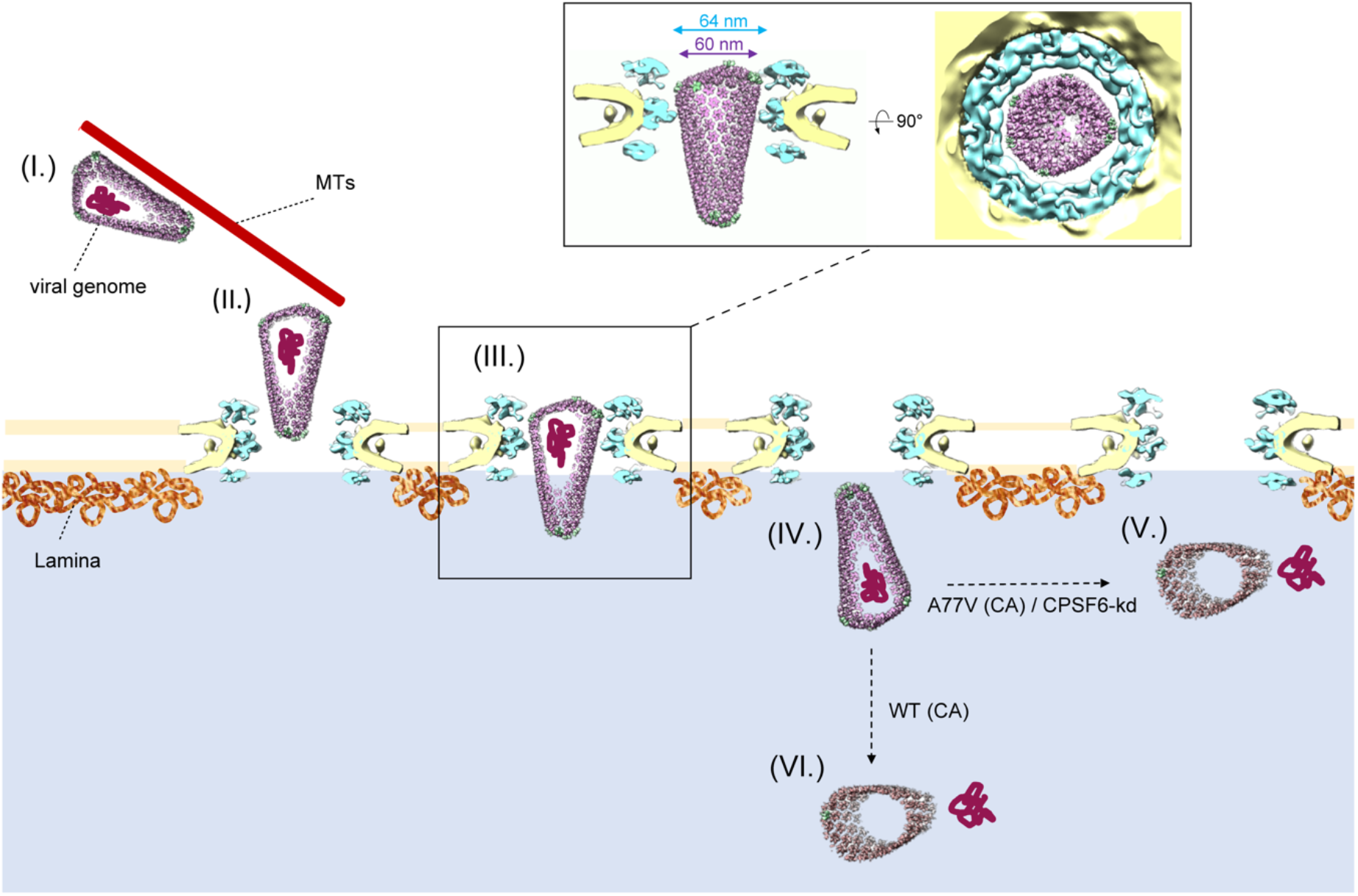
Conceptual model of HIV-1 nuclear import in T cells. In the cytosol, the intact cone-shaped HIV-1 capsid encasing the viral genome travels along microtubules towards the nuclear periphery (I.), where it docks to the NPC with the narrow CA pentamer-rich end (II.). Subsequently, the capsid penetrates into the central channel (III.). Superimposition of the mature HIV-1 capsid as determined from intact virions (Mattei et al., 2016) onto the *in cellulo* structural model of the NPC from infected cells (this study) reveals that the diameter of the NPC central channel is sufficiently wide for the transport of intact HIV-1 capsid. Side view (left) and top view (right) are shown as inset (III.). The intact HIV-1 capsid translocates into the nucleus (IV.). When CPSF6 binding is perturbed, the capsid uncoats and releases the genome at the NPC nuclear basket region (V.). When CPSF6 is available for interaction with the CA lattice, uncoating occurs deeper in the nucleoplasm (Burdick et al., 2020). In both cases (V. and VI.), the viral genome is released and integration into the host genome occurs close to the site of release. Legend: Microtubule (MTs) in red, viral genome in dark purple, lamina in orange are labeled accordingly; nuclear envelope in yellow; NPC density in cyan; intact conical HIV-1 capsid (Mattei et al., 2016) composed by hexamers in magenta and pentamers in green; uncoating of HIV-1 capsid is represented by different shape and change in color from magenta to salmon.

Although the role of microtubule-dependent transport in HIV-1 post-entry transport is well documented (Dharan et al., 2017; Fernandez et al., 2015; Malikov et al., 2015), the frequency of microtubules closely associated with intact HIV-1 capsids in direct proximity to NPCs was striking. This observation may be related to cytosolic, microtubule-associated NUP358 in addition to NPC-associated NUP358 acting as a docking station for the capsid as discussed above. NUP358 was previously suggested to relocate from NPCs into the cytosol to recruit PIC for nuclear import by recruiting CA with kinesin-1 onto microtubules (Dharan et al., 2016). During oogenesis, NUP358 condenses into granules that are actively transported along microtubules (Hampoelz et al., 2019), while in somatic cells its BicD2 binding domain mediates the association with microtubules and fulfills a role in nuclear positioning (Splinter et al., 2010). It remains to be studied whether these functions are potentially highjacked by the virus in order to utilize microtubules as platform for the delivery of HIV-1 capsids directly to their docking position at the NPC.

Because of a size mismatch of reported structures of the NPC central channel and the HIV-1 capsid, it was assumed that HIV-1 capsids need to disassemble prior to nuclear import or alternatively, NPCs are remodeled to promote nuclear entry (Monette et al., 2011). Here we demonstrate that neither needs to be the case, since the native NPC conformation in infected SupT1-R5 cells allows for the passage of intact capsids. The overall NPC architecture revealed in our analyses was not fundamentally altered, but rather dilated as compared to previous analysis of isolated nuclear envelopes (von Appen et al., 2015) that showed NPCs in a more constricted conformation. Although we cannot formally exclude that some HIV-1 particles undergo uncoating before nuclear import or during NPC translocation (Novikova et al., 2019), our data argue against it. The vast majority of CLEM ROIs analyzed in the cytosol comprised capsids containing dense material, whereas empty, tube-shaped or perturbed capsids were mainly detected in the nucleoplasm. Most importantly, the cryo-ET data presented here clearly demonstrate that intact capsids are capable of penetrating the central channel of the NPC.

While a few HIV-1 complexes at the nuclear basket of NPCs or within the nucleus still appeared conical, the majority of the identified nuclear complexes were morphologically altered and lacked interior dense material. The remnant structures detected suggest that capsid lattices are not entirely disassembled upon nuclear entry, but rather disrupted. In our CLEM experiments IN fluorescence was observed also for disrupted capsids emphasizing that at least part of the IN.mScarlet protein must stay associated with or proximate to the broken capsid at this stage. Opening the conical capsid potentially relieves strain imposed by the CA pentamers and may also be triggered by completion of reverse transcription, subsequently releasing the viral genomic cDNA. This interpretation of our observations is consistent with the finding that blockage of the nuclear pore prevents completion of reverse transcription (Dharan et al., 2020), and the concept that the generation of double-stranded DNA within the capsid may impose mechanical strain from the inside (Rankovic et al., 2017).

On a speculative note, our observations may explain the conical shape of HIV-1 capsids. The role of CA pentamers in defining capsid curvature and closure of the shell encasing the viral genome is well established (Ganser et al., 1999; Mattei et al., 2016). Beyond that, the pentamers localized towards the ends of the cone might also guide the perpendicular orientation of HIV-1 capsids with respect to NPCs during docking, and it may be hypothesized that preferential binding of the narrow ends to the NPC is mediated by the stronger enrichment of pentamers in this area of the cone. In a recent report (Lau et al., 2020), the authors established an *in vitro* system of self-assembled CA N21C/A22C (Pornillos et al., 2011) spheres, which adopted the same pentamer-hexamer and hexamer-hexamer interaction interfaces as found in the highly curved ends of the HIV capsids. They showed that CypA binds to those regions with a higher stoichiometry than to the tubular hexameric lattice and hypothesized that CA pentamers might represent specialized binding sites that are recognized by cyclophilin domains contained in host proteins, such as CypA and NUP358. While binding of CypA to the capsid had rather an inhibitory effect on HIV-1 infection and nuclear import (Burdick et al., 2017; Schaller et al., 2011), the NUP358-CA interaction at the cytoplasmic face of the NPC and potentially on microtubules is crucial for nuclear import of HIV-1 PIC (Burdick et al., 2017; Dharan et al., 2016; Schaller et al., 2011). Our data are consistent with a model in which the interaction of the narrow end of the HIV-1 capsid with the cyclophilin domain of NUP358 facilitates a capsid orientation that is advantageous for the subsequent penetration through a dense meshwork of FG-NUPs located within the central channel.

Nuclear import of the PIC was suggested to be promoted by consecutive binding of NUP153 and CPSF6 to the CA lattice (Bejarano et al., 2019). The capsid is exposed to these factors only once it penetrates deep beyond the central channel and reaches into the nuclear basket region of the NPC. Our data strongly suggest that capsid remains intact until this stage. Interestingly, the pentamers feature an open pocket for NUP153 and CPFS6 binding (Mattei et al., 2016) underscoring that binding to these factors during the late stages of nuclear import may be linked to the disruption of the lattice.

Taken together, this study uncovered the structural status of the HIV-1 capsid while it exploits the host cell transport mechanisms and protects the viral genome against detection by innate immune sensors of the cell. Our findings shift the paradigm of capsid uncoating from a total disassembly of CA proteins from the viral genome before or during translocation through the NPC to partial opening of capsids with release of viral genome occurring upon nuclear entry. This is enabled by a dilated conformation of transporting NPCs that is observed under the relevant conditions *in cellulo*.

## Acknowledgments

We thank Robert Doms, Anna Cereseto, David Bejarano and Jessica Daecke for providing cell lines or plasmid constructs. We are grateful to Maria Anders-Össwein and Vera Sonntag-Buck for expert technical assistance. We thank William Wan for support with the analysis of HIV-1 CA hexamers, Julia Mahamid for assistance at cryo-FIB microscope and Robin Burk for assistance during CLEM workflow establishment. We are grateful to Dirk Grimm for providing the AAV triple shRNA system. We thank Janina Baumbach, Kar Ho Herman Fung and Ines de Castro for valuable discussions. We would like to acknowledge the assistance of Wim Hagen and Felix Weis from the Electron Microscopy Core Facility at EMBL, Heidelberg and the assistance of Charlotta Funaya and Stefan Hillmer from Electron Microscopy Core Facility at Heidelberg University. We would like to acknowledge the support of Vibor Laketa from the Infectious Diseases Imaging Platform (IDIP) at the Center for Integrative Infectious Disease Research, Heidelberg.

The following reagents were obtained through the AIDS Reagent Program, Division of AIDS, NIAID, NIH: efavirenz from the Division of AIDS, NIAID and anti-HIV-1 p24 Hybridoma (183-H12-5C) (Cat#1513) from Dr. Bruce Chesebro. This work was funded in part by the Deutsche Forschungsgemeinschaft (DFG, German Research Foundation) – Projektnummer 240245660 – SFB 1129 project 5 (HGK), project 6 (BM) and project 20 (ML and MB) by the TTU HIV in the DZIF (HGK), EMBL and the Max Planck Society (MB).

## Author Contributions

V.Z. conceived the project, designed and performed experiments, analyzed data and wrote manuscript; E.M. conceived the project, designed and performed experiments, analyzed data and wrote manuscript; B.T. performed subtomogram averaging of CA hexamers; T.G.M. designed and performed experiments and analyzed data; C.E.Z., S.M., M.A. performed computational analysis; K.B. performed experiments and analyzed data; J.R. performed experiments and analyzed data; B.M. and M.L. supervised the project; H.-G.K. conceived the project, designed experiments, supervised the project, wrote the manuscript; M.B. conceived the project, designed experiments, supervised the project, wrote the manuscript.

## Supplementary Figures

**Figure S1.**
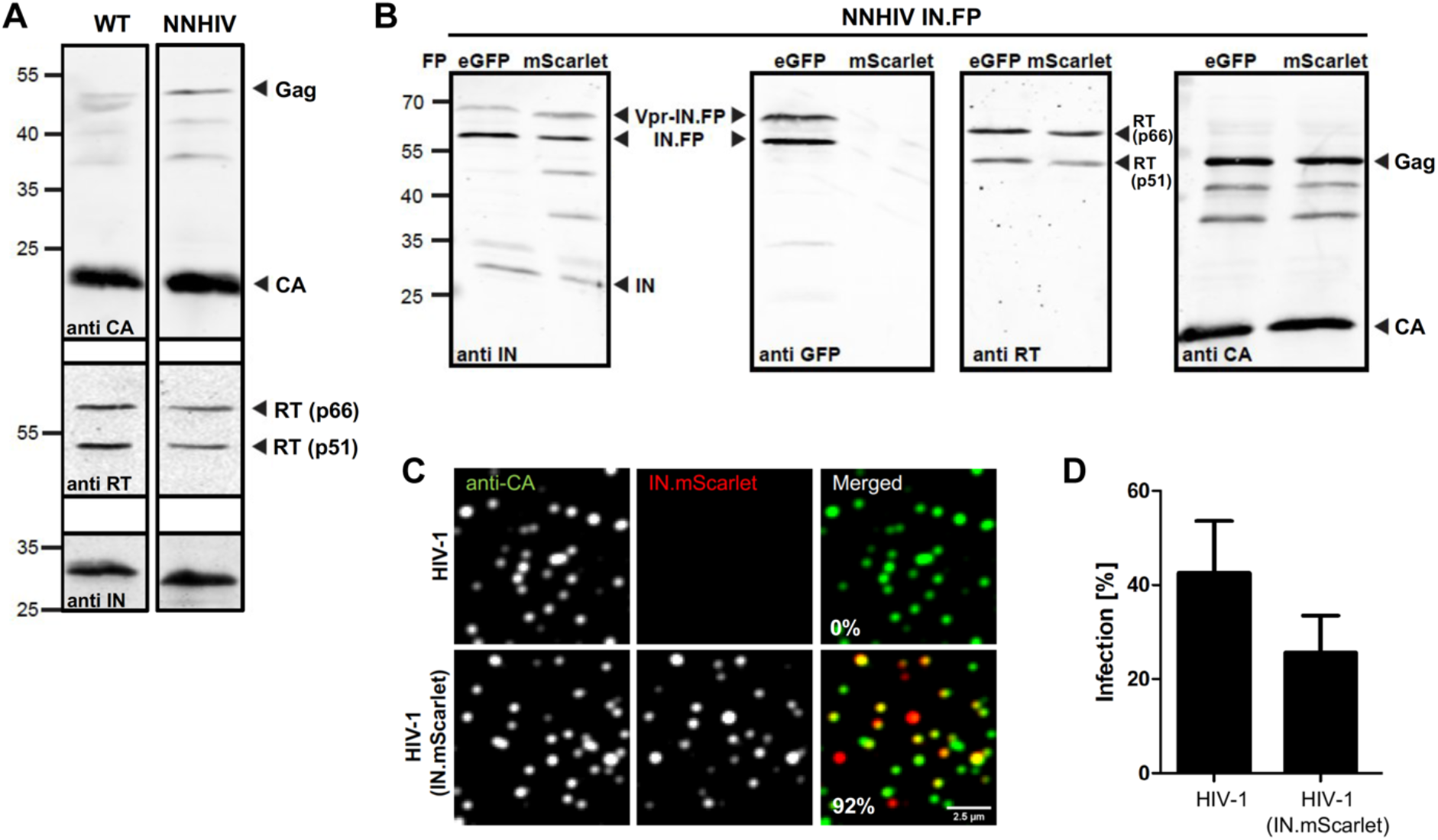
Characterization of purified NNHIV particles and the effect of IN.mScarlet incorporation on HIV-1 infectivity. **(A, B)** Immunoblot analysis of virus particles purified from the supernatant HEK 293T cells transfected with proviral plasmid pNLC4-3 (A, lane 1), pNNHIV (A, lane 2), pNNHIV and pVpr.IN_D64N/D116N_.eGFP (B, lanes 1) or pNNHIV and pVpr.IN_D64N/D116N_.mScarlet (B, lanes 2). Antisera raised against recombinant HIV-1 CA, RT or IN, or against eGFP was used. **(A)** Wildtype (WT) HIV-1 virions and NNHIV particles have a similar composition and Gag processing. **(B)** NNHIV particles labeled with IN fused to eGFP or mScarlet fluorescent protein (FP) have a comparable composition, Gag processing and FP-fused IN incorporation. **(C, D)** Effect of IN.mScarlet incorporation on HIV-1 infectivity. Particles were purified from the supernatant of HEK 293T cells transfected with plasmid pNLC4-3 (HIV-1) or pNL4-3 and pVpr.IN.mScarlet (HIV-1 IN.mScarlet). **(C)** Analysis of labeled virus particles by immunofluorescence. Virions were adhered to PEI-coated 8-well chamber glass bottom, fixed and immune-stained using antiserum against HIV-1 CA. Representative confocal images recorded in the CA (green) and IN.mScarlet (red) channel. The presented fraction (%) of labeled virions (n = 1036) was quantified using spot detector of the Icy software as described in Materials and Methods. **(D)** Infectivity of IN.mScarlet-labeled virions. SupT1-R5 cells were infected with equal amounts of non-labeled or IN.mScarlet-labeled HIV-1_NL4-3_ virions for 24 h prior to addition of T-20 fusion inhibitor. At 48 h p.i., cells were fixed and immunostained for intracellular HIV-1 CA. Levels of infected cells were scored by flow cytometry. The graph shows mean values and SEM from three independent experiments performed in triplicates.

**Figure S2.**
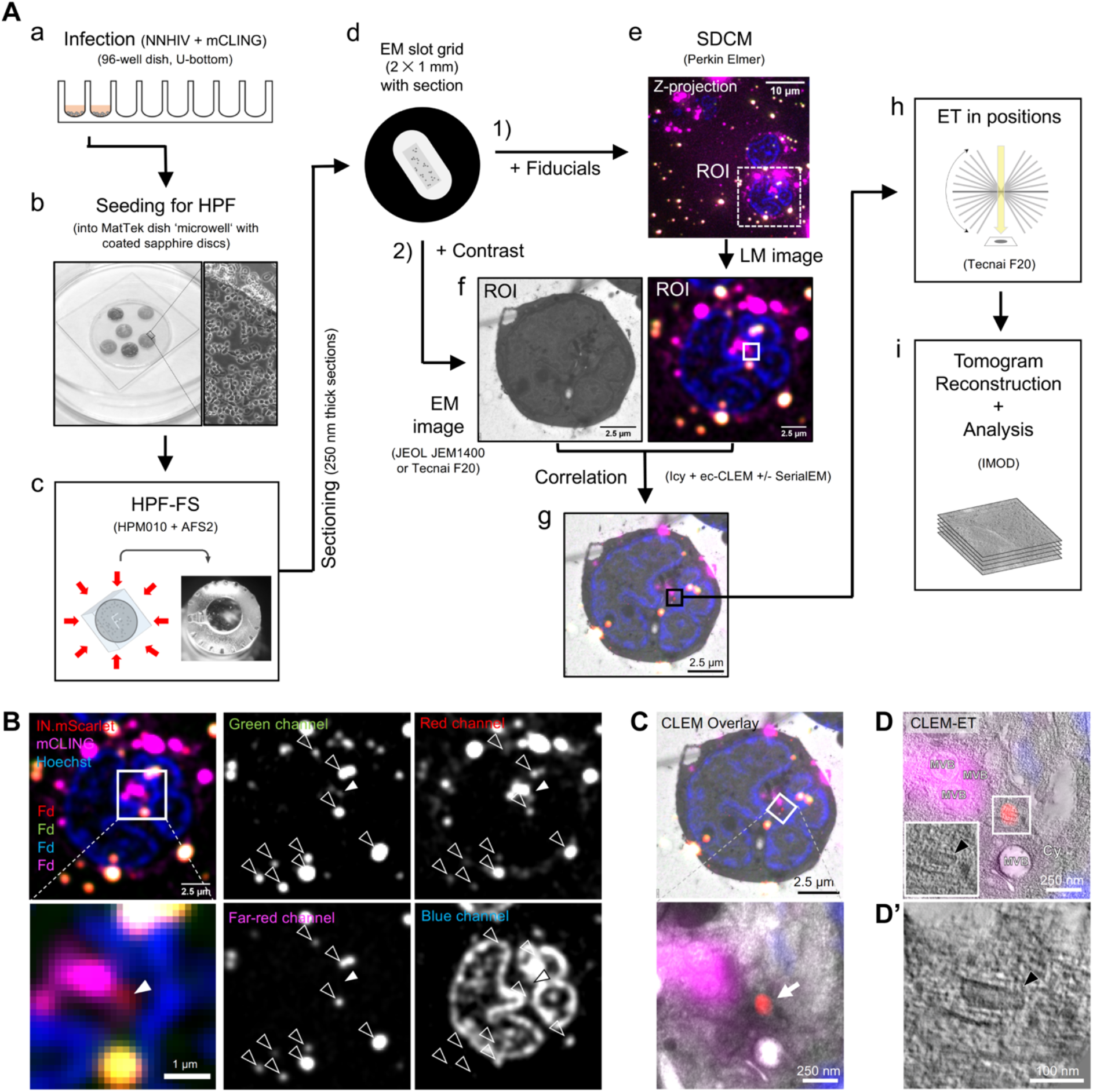
Workflow for CLEM-ET visualization of HIV-1 post-entry complexes. **(A)** SupT1-R5 cells are incubated with IN.mScarlet-carrying NNHIV particles for 90 min at 16°C in a 96-well dish. Cells are stained with mCLING.Atto647N for an additional 10 min at 16°C and shifted to 37°C to initiate virus entry (a). For high pressure freezing, cells are transferred (at 37°C, in the presence of mCLING) to MatTek dish ‘microwell’ containing carbon- and retronectin-coated sapphire discs (b). At indicated times after temperature shift, cells adhered on sapphire discs are high pressure frozen, freeze substituted and embedded in Lowicryl resin (c). After polymerization, 250 nm-thick sections of the resin-embedded cell monolayer are transferred onto EM slot grids (d). Multichannel fluorescent fiducials (TetraSpeck microspheres) are applied to the grid and examined by spinning disc confocal microscopy (SDCM) (e). Regions of interests (ROI) are identified in the resulting z-stacks (f, right) and sections on grids are contrasted for visualization by transmission electron microscopy (EM) (f, left). Using multi-fluorescent fiducials, which are visible in EM micrographs as dense 100 nm spheres; LM and EM images are correlated to identify positions of IN.mScarlet in the cell section (g). Finally, tilt series are acquired at the positions of the identified ROIs (h), tomograms are reconstructed and further analyzed (i). **(B-D’)** Identification and visualization of post-entry HIV-1 complexes in a representative resin section by 3D CLEM. **(B)** SDCM image of a 250-nm thick resin cell section. The fluorescence of IN.mScarlet (red) and mCLING.Atto647N (magenta) is retained after freeze substitution and resin embedding. Resin embedded sections are stained with Hoechst (blue) and decorated with multi-fluorescent fiducials for correlation (Fd; empty arrowheads). **(C)** CLEM overlay of a fluorescence image with the respective electron micrograph (top) and enlarged detail corresponding to ROI where a tilt series was acquired (image below). The white arrow indicates the position of the IN.mScarlet signal. **(D-D’)** Slices through tomographic reconstructions as overview correlated with SDCM image (D) and an enlarged tomographic slice (D’) show the cone-shaped NNHIV capsid in the cytosol (black arrowheads) at the position of IN.mScarlet signal. Cy, cytosol; MVB, multivesicular body.

**Figure S3.**
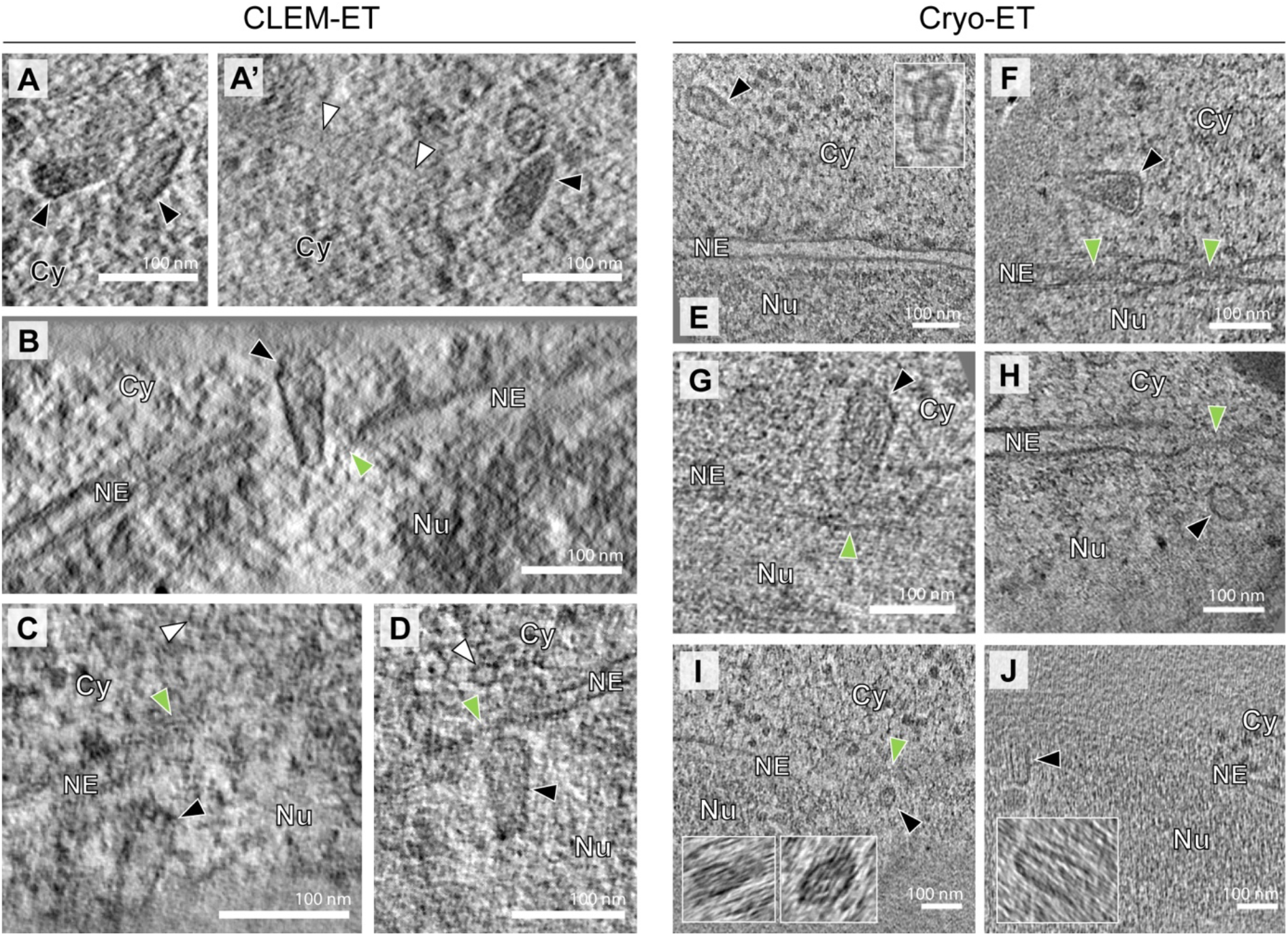
Gallery of CA A77V structures visualized at multiple stages of nuclear entry. SupT1-R5 cells were infected with IN.mScarlet carrying NNHIV-A77V CA mutant for 15 h at 37°C, prior to high pressure freezing (for CLEM) or plunge freezing (for cryo-ET). **(A-J)** Slices through tomographic reconstructions highlighting the morphology of A77V capsids or capsid-related structures visualized by CLEM-ET (A-D) or cryo-ET (E-J). Shown are examples of NNHIV-A77V structures visualized in the cytosol (A, E-G), during docking (F, G) NPC penetration (B) and after translocation through the nuclear pores (C, D, H-J). Capsids in panels (A, A’) and (B, I) were identified in the same electron tomograms. Black, white and green arrowheads indicate capsids (or capsid related structures), microtubule cross sections and NPCs, respectively. Cy, cytosol; Nu, nucleus; NE, nuclear envelope.

**Figure S4.**
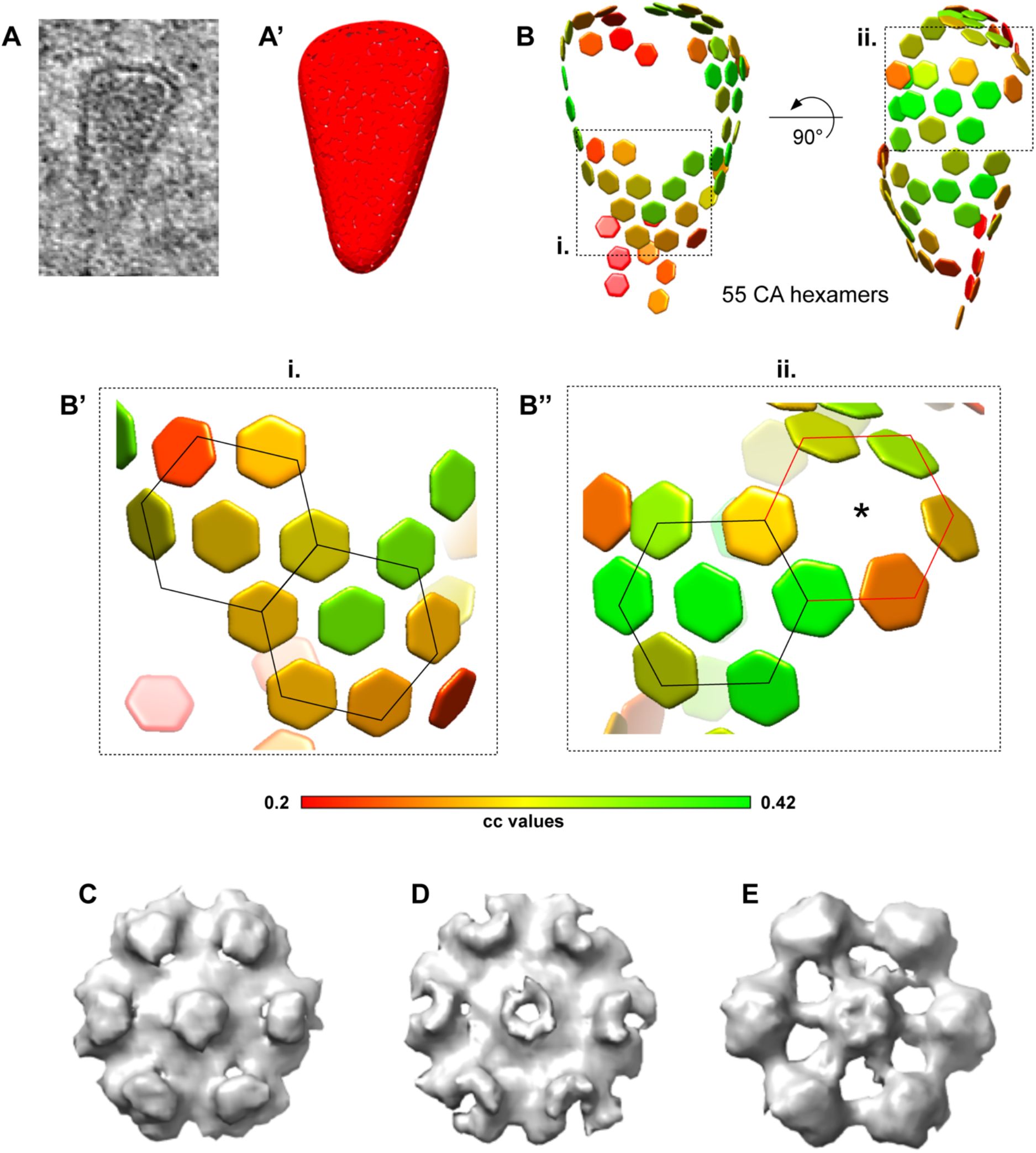
Subtomogram averaging of hexamers and reconstruction of the capsid lattice. **(A-B’’)** Distribution of hexamers along the surface of a conical HIV-1 capsid before nuclear entry. **(A)** HIV-1 capsid in the cytosol of a cryo-FIB milled SupT1-R5 cell. Corresponding tomographic slice shown in Figure S3B. **(A’)** Segmented surface for the extraction of the initial subtomograms visualized in UCSF Chimera (Pettersen et al., 2004). The following procedure was applied on the identified structures to extract subtomograms for subtomogram averaging: (i) each capsid was manually segmented in IMOD, (ii) filtered volumes were generated from each segmentation in MATLAB and (iii) randomly distributed positions were extracted with approximately 4× oversampling along the volume by defining their distance (based on the 10 nm hexamer-hexamer spacing) and the contour level of volume. **(B)** Lattice of 54 CA hexamers obtained after performing subtomogram averaging and cleaning the overlapping aligned subtomograms by cross-correlation (cc) values. cc values for each CA hexamer in the volume are shown color-coded as indicated in the colormap. **(B’-B’’)** The areas highlighted as i. and ii. of the lattice shown in (B), contain consecutive regular arrangements of six CA hexamers (highlighted by black lines) surrounding a seventh, central CA hexamer; in **(B’’)** six CA hexamers are localized to a highly curved region (red lines). The position of central CA hexamer was not detected (star). **(C-E)** Hexagonal subtomogram averages obtained from capsids in the cytosol (C) (conical capsid shown in panel B and in Figure S3B), within the NPC central channel (D) (conical capsid shown in Figure 3C), in the nucleus (E) (tubular capsid shown in Figure 4D).

**Figure S5.**
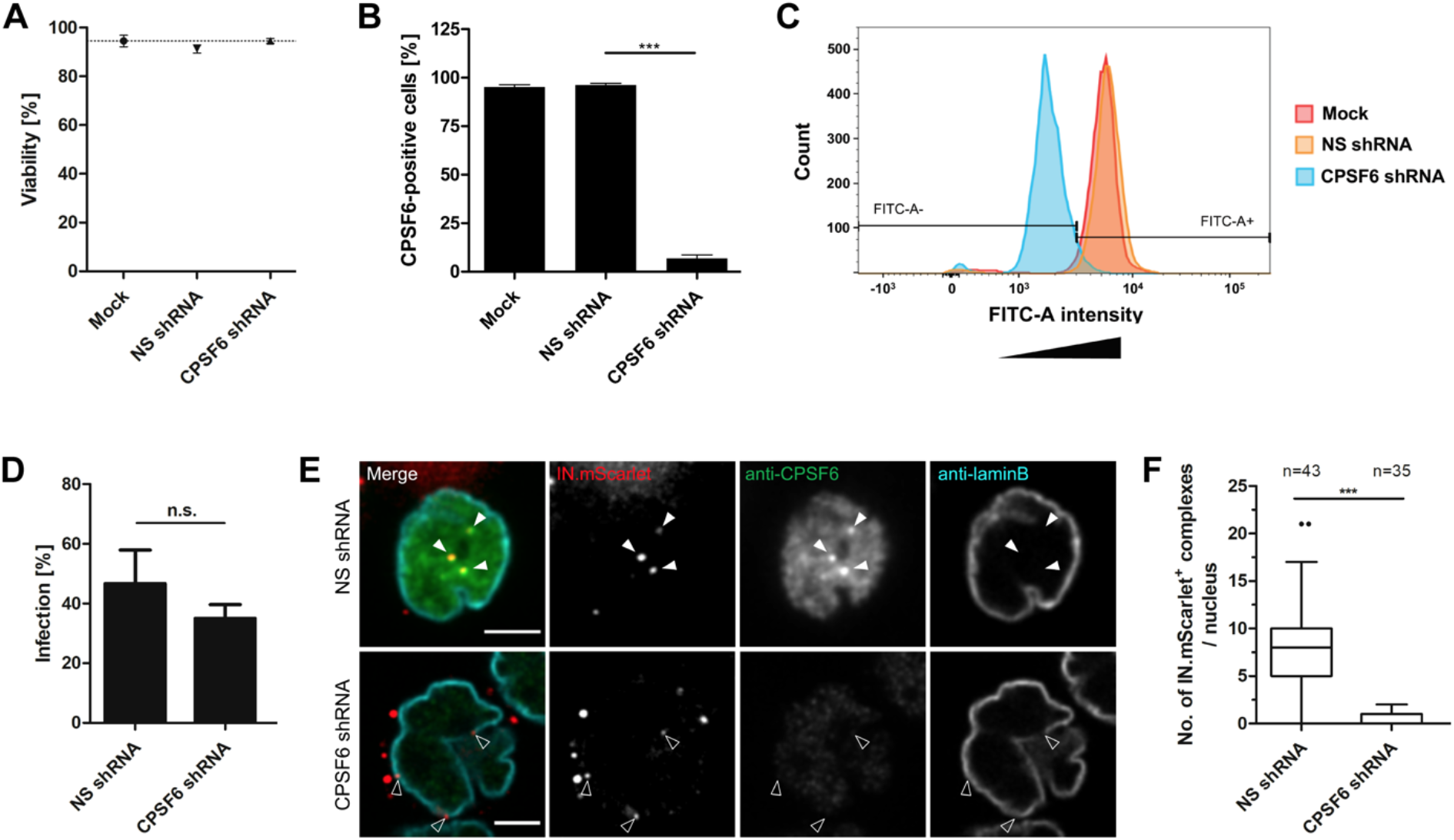
HIV-1 infectivity and nuclear entry upon CPSF6 knock-down in SupT1-R5 cells. **(A-C)** CPSF6 knock-down in SupT1-R5 cells. Cells were transduced with AAVs expressing non-silencing (NS) or CPSF6 shRNA. Mock-transduced cells were used as additional control. 72 h after transduction, cell viability and CPSF6 knock-down efficiency were analyzed. **(A)** The number of viable cells was determined by trypan blue exclusion. The graph shows mean values and SD from three independent experiments. **(B)** CPSF6 knock-down efficiency. Cells were fixed, immunostained with anti-CPSF6 antibody, followed by staining with secondary antibody conjugated with AlexaFluor 488. The proportion of CPSF6-positive cells was determined by flow cytometry. Data are representative of three independent experiments. Statistical significance was calculated using an unpaired two-tailed t-test, *** p < 0.0001. **(C)** Representative histogram of CPSF6 signal intensity for cell populations analyzed by FACS. For the experiment shown, 12,332 (mock), 15,195 (NS shRNA) and 13,454 (CPSF6 shRNA) events were analyzed. **(D-F)** HIV-1 infectivity and nuclear import in SupT1-R5 cells upon CPSF6 knock-down. **(D)** Effect of CPSF6 knock-down on HIV-1 infectivity. AAV-transduced cells were infected with wild-type HIV-1_NL4-3_. At 48 h p.i., cells were fixed, immunostained for intracellular HIV-1 CA and infection was scored by flow cytometry. The graph shows mean values and SEM from three independent experiments performed in triplicates. Statistical significance was assessed by unpaired two-tailed Student’s t test. n.s., not significant. **(E, F)** Effect of CPSF6 knock-down on virus nuclear entry. AAV-transduced cells were infected with IN.mScarlet carrying NNHIV particles for 90 min at 16°C and then shifted to 37°C to initiate virus entry. Cells were fixed 15 h after temperature shift and immunostained for CPSF6 (green) and lamin B (cyan). Arrowheads indicate IN.mScarlet-positive complexes (red) in the nucleus (white arrowheads) of cell expressing non-silencing (NS) control shRNA, and IN.mScarlet signals associated with nuclear envelope (empty arrowheads) in cell expressing CPSF6 shRNA. Scale bars: 2.5 μm. **(F)** Box-and-whisker plot shows the numbers of IN.mScarlet positive complexes in nuclei of cells expressing NS or CPSF6 shRNA. Outliers were identified by Tukey’s test. Statistical significance was assessed by unpaired two-tailed Student’s t test. ***: p<0.0001.

**Figure S6.**
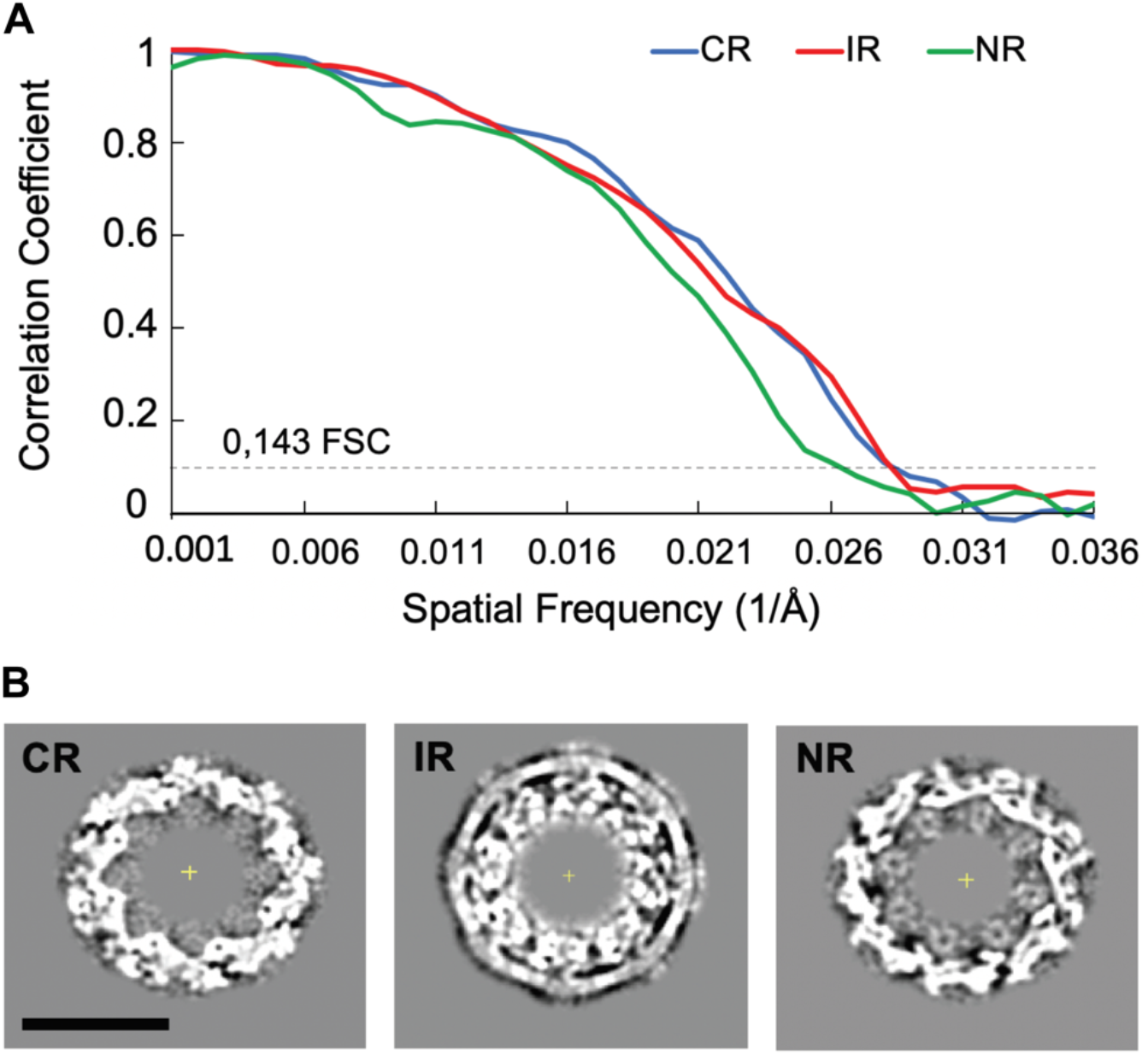
Cryo-EM structure of actively transporting NPC upon HIV-1 infection. **(A)** Resolution estimation by gold standard Fourier shell correlation (FSC) for the cryo-EM map of the NPC from NNHIV infected SupT1-R5 cells. After alignment of the whole pore, each NPC was splitted into the even and odd subunits. The cytoplasmic (CR), nuclear (NR) and inner ring (IR) from each subunit were independently subjected to subtomogram averaging. The gold standard FSC was calculated with FSC server - EMBL-EBI with half maps as input. The curves for CR, NR and IR intersect the 0.143 criterium respectively at the resolution of 36.3 Å, 35.8 Å, and 39.9 Å. **(B)** Slices through the maps at the level of the cytoplasmic ring (CR), inner ring (IR) and nuclear ring (NR). Scale bar: 50 nm.

**Video S1: Morphology of HIV-1 complexes in the cytosol.** Related to Figure 2D. Tomographic reconstruction and isosurface rendering highlights a microtubule (MT)-associated cone-shaped NNHIV capsid located in proximity of NPC. MT red; capsid, magenta; NE, yellow; NPC, cyan (this study).

**Video S2: Morphology of HIV-1 complexes docking to the NPC.** Related to Figure 2E. Tomographic reconstruction and isosurface rendering highlighting a microtubule (MT)-associated conical NNHIV capsid docking to the NPC. MT red; capsid, magenta; NE, yellow; NPC, cyan (this study).

**Video S3: Morphology of HIV-1 complexes inside of the nucleus.** Related to Figure 2F. Tomographic reconstruction showing the morphology of clustered, capsid-related NNHIV structures within the nucleus.

**Video S4: Intact CA-A77V capsids penetrate into the NPC central channel.** Related to Figure 3A. Tomographic reconstruction showing visually intact NNHIV A77V capsid deep inside of the central channel of the NPC, exposing its narrow end towards the nucleoplasm.

**Video S5: Cone-shaped CA-A77V capsids enter the NPC central channel.** Related to Figure 3B. The movie shows orthoslices through a representative cryo-electron tomogram and an isosurface-rendered view of a CA-A77V capsid (magenta), the nuclear envelope (yellow) and the associated NPC (cyan; this study).

**Video S6: Morphologically altered CA-A77V capsids are observed in close proximity to the NPC in the nucleoplasm.** Related to Figure 4D. The movie shows orthoslices through a representative cryo-electron tomogram and an isosurface-rendered view of MTs (red), a CA-A77V capsid (magenta), the nuclear envelope (yellow) and the associated NPC (cyan; this study).

**Video S7: Morphology of NPC-associated HIV-1 complexes upon CPSF6 knock-down.** Related to Figure 5C. Tomographic reconstruction showing an empty capsid-related NNHIV structure retained at the nucleoplasmic side of the NPC in SupT1-R5 cell upon CPSF6 knockdown.

## Materials and Methods

### Cell cultures

Human T lymphoblast cells SupT1-R5 (stably expressing exogenous CCR5 under puromycin selection; a kind gift from Robert Doms, University of Pennsylvania, USA; certified by Eurofins according to DAkkS ISO 9001:2008) were cultivated at 37°C in a humidified incubator with a 5% CO_2_ atmosphere, using RPMI 1640 medium with GlutaMAX (ThermoFisher Scientific) supplemented with 10% fetal bovine serum (FBS; Merck), 50 U/ml of penicillin, 50 μg/ml of streptomycin (ThermoFisher Scientific) and 0.3 μg/ml puromycin (Merck). Human embryonic kidney 293T cells (HEK 293T) (Pear et al., 1993) were maintained in Dulbecco’s modified Eagle medium (DMEM; ThermoFisher Scientific) supplemented with FBS, penicillin and streptomycin at concentrations as above.

### Plasmids

Plasmid pNLC4-3 for the production of infectious HIV-1 particles was described previously (Bohne and Kräusslich, 2004). Plasmid pNNHIV (pNLC4-3 IN_D64N/D116N_ tat_Δ33-64bp_) for production of non-infectious, RT-competent HIV-1 particles, was constructed by successive introduction of integrase catalytic mutations D64N and D116N by Quickchange PCR into pUC19 NL4-3_5’CA-3’Vpr_ (generated by subcloning a 4296 bp SphI/EcoRI fragment from pNL4-3 into pUC19, from which the NdeI site has been removed by Klenow insertion and religation). For mutagenesis, following primers were used: IN_D64N_ forward, 5’-GTAGCCCAGGAATATGGCAGCTAAACTGTACACATTTAGAAGGAAAAG-3’; IN_D64N_ reverse, 5’-CTTTTCCTTCTAAATGTGTACAGTTTAGCTGCCATATTCCTGGGCTAC-3’; IN_D116n_ forward, 5’-GCAGGAAGATGGCCAGTAAAAACAGTACATACAAATAATGGCAGCAATTTCACCAGTACTACAGTTAAGG-3’; IN_D116N_ reverse, 5’-CCTTAACTGTAGTACTGGTGAAATTGCTGCCATTATTTGTATGTACTGTTTTTACTGGCCATCTTCCTGC-3’). Afterwards, a fragment containing the IN mutations was subcloned into the pNLC4-3 tatΔ33-64bp backbone which contains a 31 bp deletion in the first exon of tat (Bejarano et al., 2019) using AgeI/EcoRI creating pNNHIV. Plasmid pNNHIV-A77V carrying an A77V mutation in the CA-coding region of *gag* was constructed through double digestion of pNL4-3-A77V (Bejarano et al., 2019) with BssHII and AgeI, followed by ligation with the corresponding fragment from pNNHIV backbone. Plasmid pVpr.IN.mScarlet encoding a Vpr.IN.mScarlet fusion protein with a HIV-1 protease recognition site between Vpr and IN was constructed from pVpr.IN.eGFP (Albanese et al., 2008) by PCR amplification of the mScarlet gene from pmScarlet C1 (Addgene Cat#85042; (Bindels et al., 2017)) (primers used for PCR: forward, 5’-AGGACGAGGACCGGGATCCACCGGTCGCCACCATGGTG-3’; reverse, 5’-TGATTATGATCTAGAGTCGCTTACTTGTACAGCTCGTCCATGCC-3’) and cloning into AgeI/NotI linearized pVpr.IN.eGFP substituting eGFP for mScarlet using Gibson assembly. Plasmid pVpr.IN_NN_.mScarlet (Vpr.IN_D64N/D116N_.mScarlet) for labeling of non-infectious NNHIV particles was constructed similarly by using pVpr.IN_D64N/D116N_.eGFP (gift from D. A. Bejarano) as backbone.

Plasmids for AAV production, AAV helper plasmid encoding *rep* and 1P5 *cap* gene (Börner et al., 2020), the vector for AAV expression of triple short hairpin RNA (shRNA) targeting three CPSF6 sequences (Bejarano et al., 2019) and vector for expression of a single non-silencing shRNA (Börner et al., 2010), and adenoviral helper plasmid providing helper functions for AAV production (Matsushita et al., 1998) were described previously.

### Antibodies and reagents

For immunofluorescence staining, rabbit polyclonal antiserum against HIV-1 CA raised against purified recombinant protein (in house) (Welker et al., 2000), mouse monoclonal antibody against lamin B (sc-365962; Santa Cruz) and affinity-purified rabbit polyclonal antibody against human CPSF6 (HPA039973; Merck) were used at a dilution of 1:1,000, 1:200 and 1:250, respectively. Secondary antibodies donkey anti-rabbit IgG and donkey anti-mouse IgG conjugated with Alexa Fluor 488 and 647, respectively (all purchased from ThermoFisher Scientific), were used at 1:1,000 dilution. For western blot analyses, we used antisera raised against purified recombinant proteins (in house): sheep polyclonal antiserum against HIV-1 CA (Müller et al., 2009), rabbit polyclonal serum against HIV-1 IN (Welker et al., 2000), rabbit polyclonal serum against HIV-1 RT (Müller et al., 2004) and rabbit polyclonal antiserum against GFP (Müller et al., 2004). Sera were used at a dilution 1:5,000, 1:1,000, 1:1,000 and 1:1,000, respectively. For detection of HIV-1 CA by flow cytometry, fluorescein isothiocyanate (FITC)-conjugated mouse monoclonal antibody KC57 (Beckman Coulter) was used at a dilution of 1:100.

A stock solution of 50 μM mCLING.Atto647N (710006AT1; Synaptic Systems, Göttingen, Germany) was prepared in PBS (pH 7.4) was stored at −80°C. A stock solution of 10 mM efavirenz (obtained through the AIDS Research and Reference Reagent Program, Division AIDS, NIAID) was prepared in dimethyl sulfoxide and stored at −20°C. A 20 mM stock solution of T-20 (enfuvirtide; Roche) was prepared in H_2_O and stored at −20°C. A 16 μg/ml solution of Retronectin (T100B; Takara Bio Inc.) was prepared in PBS and stored at −20 °C. Retronectin was recycled up to 7 times.

### Virus and virus-like particles

To produce infectious HIV-1 virions (HIV-1_NL4-3_) (Adachi et al., 1986) or RT-competent viruslike particles (NNHIV), HEK 293T cells grown on 175 cm^2^ tissue culture flasks side bottom were transfected with pNLC4-3 or pNNHIV (both, 70 μg DNA per flask), using calcium phosphate precipitation according to standard procedures. For production of infectious virions labeled with IN.mScarlet, cells were co-transfected with pNLC4-3 and pVpr.IN.mScarlet at a molar ratio of 4.5:1. To produce NNHIV particles or their A77V CA mutation-carrying version labeled with IN.mScarlet, cells were co-transfected with pNNHIV or pNNHIV-A77V and pVpr.IN_NN_.mScarlet at a molar ratio of 4.5:1. Culture media from virus-producing cells were harvested at 44–48 h post-transfection, cleared by filtration through a 0.45 μm nitrocellulose filter, and particles from media were concentrated by ultracentrifugation through a 20% (w/w) sucrose cushion at 90,000 × g for 90 min at 4°C. Particles were resuspended in PBS containing 10% FBS and 10 mM HEPES (pH 7.2) and stored in aliquots at −80°C. For detection of HIV-1 RT products by ddPCR, virus-containing medium from producing cells was treated with 15 U/ml DNase I (Merck) and 10 mM MgCl_2_ for 5 h at 37°C prior to ultracentrifugation. Particles were then aliquoted and stored as above. Particle-associated RT activity was determined by SG-PERT (SYBR Green-based Product-Enhanced Reverse Transcription assay) (Pizzato et al., 2009).

To produce AAV vectors, HEK 293T cells grown on 15-cm dishes were transfected using polyethylenimine (PEI) (Merck), a transfection reagent for standard triple transfection protocol. Cells were co-transfected with AAV helper plasmid encoding AAV *rep* and 1P5 *cap* gene, an AAV vector plasmid for expression of shRNA and adenoviral helper plasmid at a molar ratio of 1:1:1. At 72 h after transfection, cells were collected in PBS followed by centrifugation at 500 × g, 15 min, at room temperature. The cell pellet was resuspended in 20 ml Benzonase buffer, lysed by freeze-thaw cycles in liquid nitrogen and subsequent sonification. Cell debris was removed by two centrifugations, each 4000 × g, 15 min, at 4°C. The virus-containing lysate was purified via iodixanol density-gradient ultracentrifugation (290,000 × g, 120 min, at 4°C) and rebuffered in PBS using amicon spin columns (Merck). Particles in PBS were stored in aliquots at −80°C. Viral genome titers were determined by quantitative real-time PCR, using set of primers/probe for a short non-coding sequence from eGFP gene in CPSF6 multiplexing construct (forward, 5’-GAGCGCACCATCTTCTTCAAG-3’ (Michler et al., 2016); reverse, 5’-TGTCGCCCTCGAACTTCAC-3’ (Michler et al., 2016); probe, 5’-6-carboxy-fluorescein [FAM]-ACGACGGCAACTACA-black hole quencher 1 [BHQ1]-3’ (Michler et al., 2016)).

### Western Blot

Virus particles were subjected to 17.5% SDS-PAGE (200:1 acrylamide:bis-acrylamide) for 1 h at 43 mA. Proteins were transferred to a nitrocellulose membrane (GE Healthcare) by semi-dry blotting for 1 h at 0.8 mA/cm^2^. Selected viral antigens were stained with indicated primary antisera raised against purified recombinant proteins, followed by staining with corresponding secondary antibodies (LiCor). Detection was performed using a LiCor Odyssey infrared scanner according to the manufacturers’ instructions.

### Detection of HIV-1 RT products by ddPCR

Preparation of samples for detection of NNHIV RT products by digital droplet PCR (ddPCR) was performed as described in Zila et al (Zila et al., 2019). Briefly, SupT1-R5 cells were distributed in 96-well plates (3 × 10^5^ cells/well; U-bottom; Greiner Bio-One), infected with NNHIV particles (5.6 μUnits of RT/cell) and further incubated at 37°C. Cells infected with in the presence of reverse transcription inhibitor efavirenz (EFV) were used as control. At selected times post-infection, cells were washed with PBS, lyzed, proteinase K in samples was inactivated and samples were stored at −20°C. For ddPCR (Hindson et al., 2011; Morón-López et al., 2017), set of primers/probe annealing to the gag open reading frame were used to detect late RT products (forward, 5’-CATGTTTTCAGCATTATCAGAAGGA-3’; reverse, 5’-TGCTTGATGTCCCCCCACT-3’; probe, 5’-FAM-CCACCCCACAAGATTTAAACACCATGCTAA-BHQ1-3’ (Palmer et al., 2003)). 2-LTR circles were detected with another set of primers/probe (forward, 5’-CTAACTAGGGAACCCACTGCT-3’; reverse, 5’-GTAGTTCTGCCAATCAGGGAA-3’; probe, 5’-FAM-AGCCTCAATAAAGCTTGCCTTGAGTGC-BHQ1-3’ (Puertas et al., 2018)). To normalize copy numbers of HIV-1 derived DNA to the copy numbers of the housekeeping gene, the single-copy host gene encoding RNase P protein subunit p30 (RPP30) was quantified (forward, 5’-GATTTGGACCTGCGAGCG-3’; reverse, 5’-GCGGCTGTCTCCACAAGT-3’; probe, 5’-FAM-CTGACCTGAAGGCTCT-BHQ1-3’ (Hindson et al., 2011)). Preparation of reaction mixtures, droplets generation, PCR amplification and data analysis were performed as described previously (Zila et al., 2019).

### CPSF6 knock-down

CPSF6 knock-down was performed using AAV vectors. SupT1-R5 cells were distributed into 96-well plates (5 × 10^4^ cells/well; Flat bottom; Greiner Bio-One) and transduced once with equal amounts of purified AAV (2.5 × 10^6^ vector genomes/cell) expressing three shRNAs against CPSF6 or a non-targeted shRNA (NS control). Mock-transduced cells were used as an additional control. At 72 h after transduction, viability of cells was assessed by trypan blue exclusion using the TC20 Automated Cell Counter (Bio-Rad). To determine CPSF6 knockdown efficiency, cells were fixed with 4% formaldehyde (FA) in PBS and immunostained with anti-CPSF6 antibody, followed by staining with secondary antibody conjugated with Alexa Fluor 488. Proportion of CPSF6-positive cells was scored by flow cytometry, using a BD FACSCelesta flow cytometer (BD Biosciences) and data were processed using Flowjo software (FlowJo LLC, BD Biosciences).

### Infectivity assays

To determine the effect of IN.mScarlet incorporation on virus infectivity, SupT1-R5 cells were distributed into 96-well plates (3 × 10^5^ cells/well; U-bottom; Greiner Bio-One, 650180) and infected with equal amounts of non-labeled or IN.mScarlet carrying wild-type HIV-1_NL4-3_ virions (both at 1 μUnits of RT/cell). To determine HIV-1 infectivity upon CPSF6 downregulation, SupT1-R5 cells with AAV (expressing CPSF6 shRNA or non-silencing control shRNA) were distributed at 72 h post-transduction into 96-well plates (1 × 10^5^ cells/well; U-bottom; Greiner Bio-One) and infected with wild-type HIV-1 (1 μUnits of RT/cell). Infected cells were incubated for 24 h at 37°C before addition of T-20 (50 μM) fusion inhibitor to prevent a second round of infection. Infectivity was scored at 48 h p.i. by flow cytometry. For this, cells were fixed with 4% FA in PBS for 90 min at room temperature and immunostained for intracellular HIV-1 CA for 30 min at 4°C using KC57-FITC antibody (Beckman Coulter) diluted in 0.1% Triton X-100, 0.1 mg/ml BSA in PBS. Cells were analyzed using a BD FACSCelesta flow cytometer (BD Biosciences).

### Immunofluorescence Staining

To determine efficiency of HIV-1 nuclear import after CPSF6 knock-down (72 h after transduction with AAV expressing CPSF6 shRNA or non-silencing control shRNA), cells distributed in 96-wells were infected with IN.mScarlet-labeled NNHIV (5.75 μUnits of RT/cell). Cells were incubated with the virus for 14 h at 37°C, then transferred to PEI-coated wells of glass-bottom 8-well chamber slides (Ibidi, 80827) and let to adhere to the coated glass bottom for an additional 1 h at 37°C. Subsequently, cells were fixed with 4% FA in PBS (15 min at room temperature), permeabilized with 0.5% Triton X-100 (5–10 min) in PBS, washed with PBS and blocked for 30 min with 2% BSA in PBS. Immunostaining with primary and secondary antibody was carried out for 1 h each. For immunofluorescence analysis of HIV-1 particles labeled with IN.mScarlet, particles were diluted in culture media and let to adhere to PEI-coated wells of glass-bottom 8-well chamber slides (Ibidi, 80827) for 30 min at room temperature in the dark. Subsequently, samples were fixed with 4% FA in PBS, permeabilized with 0.5% Triton X-100 in PBS (5 min) and immunostained as above.

### Confocal Microscopy and Image analysis

Multichannel z-series of HIV-1 infected cells or glass bottom-adhered particles were acquired by a Nikon Ti PerkinElmer UltraVIEW VoX 3D spinning-disc confocal microscope (Perkin Elmer, Waltham, MA) using a 100× oil immersion objective (numeric aperture [NA], 1.49; Nikon), with a z-spacing of 200 nm and excitation with the 488-, 561-, and 633-nm laser lines. To quantify the efficiency of HIV-1 nuclear import after CPSF6 knock-down, a 3D volume of infected cells was reconstructed from acquired z-stacks using Imaris software (Oxford Instruments). The background was subtracted and individual IN.mScarlet signals were automatically detected using the spot detector Imaris module. Objects localized within the nucleus which were negative in the lamin B channel were classified as intranuclear. To determine the efficiency of labeling of HIV-1 virions with IN.mScarlet, the local background in the images of glass-bottom adhered virions was subtracted and CA and IN.mScarlet spots were detected using the spot detector function of the software Icy (de Chaumont et al., 2012).

The threshold was determined as the mean value of more than 200 segmented positions in the mScarlet channel where no particle was detected. Camera background subtraction, illumination correction, gauss filtering and cropping were performed using Fiji software (Schindelin et al., 2012).

### Sample preparation for CLEM

SupT1-R5 cells were distributed in well of 96-well plate (4 × 10^5^ cells/well; U-bottom; Greiner Bio-One, 650180), pelleted (3 min / 200 × g) and resuspended in complete RPMI medium containing IN.mScarlet-labeled NNHIV or NNHIV-A77V particles (25 μUnits of RT/cell, corresponding to MOI of 2.5–5 using the same μUnits of RT/cell by infectivity assay). Culture medium was supplemented with 20 mM HEPES (pH 7.2). Cells were incubated with particles for 90 min at 16°C to synchronize virus entry. For the detection of post-fusion HIV-1 complexes in the cytosol, mCLING.Atto647N (Synaptic Systems, Germany) was added after adsorption at a final concentration of 2 μM. Cells were incubated for an additional 10 min at 16°C and then shifted to 37°C to initiate virus entry. 1 h prior to high pressure freezing (HPF), cells were suspended in pre-warmed (37°C) medium supplemented with mCLING.Atto647N (2 μM) and transferred to glass-bottomed ‘microwell’ of MatTek dish (MatTek, USA) containing carbon-coated and retronectin-coated sapphire discs (Engineering Office M. Wohlwend, Switzerland). Additional coating with retronectin (Takara Bio) (16 μg/ml, overnight at 37°C) was performed to keep cells firmly attached to the sapphire disc surface during subsequent processing of the sample for high pressure freezing (HPF). For HPF, the discs were removed from the medium, placed between hexadecane-treated aluminium specimen carriers, one flat, the second with a 0.1 mm cavity and the assembled sandwich was transferred to HPM010 (Abra Fluid, Widnau, Switzerland) holder and processed by HPF. Alternatively, HPF was performed using Leica EM ICE high pressure freezer. Sapphire discs were subsequently transferred from liquid nitrogen to the freeze-substitution (FS) medium (0.1% uranyl acetate, 2.3% methanol and 1–3% H_2_O in Acetone) tempered at −90°C in FS device (Leica AFS2) equipped with a robotic solution handler (Leica FSP). Samples were FS-processed and embedded in Lowicryl HM20 resin (Polysciences, USA) according to a modified protocol of Kukulski et al (Kukulski et al., 2011): the temperature was raised to −45°C (7.5°C / h), the samples were washed four times with acetone (25 min each) and infiltrated with increasing concentrations (25, 50 and 75%; 3 h each) of Lowicryl in acetone, while the temperature was further raised to −25°C (3.3°C / h). Subsequently, the acetone-resin mixture was replaced by pure Lowicryl (for 1 h), the resin was then exchanged three times (for 3, 5 and 12 h) and samples were UV polymerized for 24 h at - 25°C. The temperature was then raised to 20°C (3.7°C / h) and UV polymerization continued for additional 24 h.

### CLEM and Electron tomography

Thick resin sections (250 nm) were obtained using a microtome (Leica EM UC7) and placed on a slot (1 × 2 mm) EM copper grids covered with a formvar film (Electron Microscopy Sciences, FF2010-Cu). Grids were placed (section face-down) for 10 min on 20 μl drops of 1× PHEM buffer (pH 6.9) containing 0.1 μm TetraSpeck beads (1:25) (ThermoFisher Scientific) as fiducial marker and 10 μg/ml Hoechst33258 () to stain nuclear regions in cell sections. Unbound or loosely attached fiducials were washed out on several drops of water and grids were transferred on 25 mm glass coverslip, which were mounted in a water-filled ring holder (Attofluor cell chamber, ThermoFisher Scientific). Z-stacks of sections were acquired by PerkinElmer UltraVIEW VoX 3D Spinning-disc Confocal Microscope (Perkin Elmer, Waltham, MA) using a 100× oil immersion objective (NA 1.49; Nikon), with a *z*-spacing of 200 nm and excitation with the 405, 488, 561 and 633 nm laser line. To identify intracellular mCLING-negative signals of IN.mScarlet, the acquired z-stacks were visually examined using Fiji software (Schindelin et al., 2012). EM grids were subsequently decorated with 15 nm protein-A gold particles on both sides for tomogram alignment and stained with 3% uranyl acetate (in 70% methanol) and lead citrate. Individual grids were then placed in a high-tilt holder (Fischione Model 2040) and loaded to a Tecnai TF20 (FEI) electron microscope (operated at 200 kV) equipped with a field emission gun and a 4K by 4K pixel Eagle CCD camera (FEI). A full grid map was acquired using SerialEM to map all cell sections (Mastronarde, 2005). To identify positions of IN.mScarlet signals for electron tomography, EM images of selected cell sections were acquired and pre-correlated with imported SDCM images of the same cell section in SerialEM using the fiducials as landmark points (Schorb et al., 2017). Single- or dual-axis electron tomograms in correlated positions were carried out. Tomographic tilt ranges were typically from −60° to 60° with an angular increment of 1°. The pixel size was 1.13 nm. Alignments and 3D reconstructions of tomograms were done with IMOD software (Kremer et al., 1996). High precision post-correlation was performed using eC-CLEM plugin (Paul-Gilloteaux et al., 2017) in Icy software (de Chaumont et al., 2012). Alternatively, EM images of selected ROIs were acquired using JEOL JEM1400 electron microscope (JEOL Ltd., Japan) operated at 120 kV and equipped with a bottom mounted 4K by 4K pixel digital camera (F416; TVIPS GmbH, Germany). Subsequent high precision correlation and electron tomography were performed as above.

### Cell vitrification and cryo-FIB milling

SupT1-R5 cells were distributed into a 96-well plate (4 × 10^5^ cells/well; U-bottom), incubated with NNHIV-A77V particles (for both, 25 μUnits of RT/cell) and vitrified at 15 h post-infection by plunge freezing as follows: 3.5 μl of cell suspension from 80-100 μl of final volume were applied on glow discharged 200-mesh EM copper grids coated with R 2/1 holey SiO_2_ films (Quantifoil Micro Tool GmbH) and plunge frozen in liquid ethane at ~−184°C using a Leica EM GP grid plunger. In the blotting chamber (maintained at 37°C temperature and 90% humidity), the grids were blotted for 2–3 sec with a filter blotting paper (Whatman 597) applied to the reverse side. The frozen grids were subsequently fixed into modified auto-grids containing a cut out to allow milling at a shallow angle (Rigort et al., 2012) and transferred to the Aquilos FIB-SEM (ThermoFisher Scientific) at liquid nitrogen temperature. Samples were sputter-coated with inorganic platinum (10 mA, 10–20 sec) and coated with organometallic protective platinum layer using the in-situ gas injection system (GIS) (Hayles et al., 2007) (7–12 sec). Ablation of undesired cellular material was performed at stage tilt angles of 18°–20° by focusing Gallium ion beam at 30 kV on parallel rectangular patterns above and below the area of interest. Lamella preparation is conducted in a stepwise milling to produce 150–250 nm sections of the biological sample. 0.3 μm gaps at 4–5 μm distance alongside the lamellae were often generated on grids to reduce lamella bending happening during the final milling step (Wolff et al., 2019). Before unloading, grids were sputter-coated again for 2 sec (10 mA) to improve conductivity of final lamellae.

### Acquisition and processing of cryo tilt series

Grids with lamellae were loaded into the Titan Krios cassette. During sample loading, the autogrids were mounted in the cassette to orient the FIB-milled lamellae perpendicularly to the tilt-axis of the microscope. Tilt series were acquired on a Titan Krios (Thermofisher Scientific, FEI) operating at 300 kV equipped with a Gatan Quantum post-column energy filter and a Gatan K2 4K by 4K direct electron detector. Prior to tilt series acquisition, an initial overview of the entire grid was acquired at 1500× magnification for the lamellae identification and subsequently each lamella on grid was mapped at 6500× magnification to identify positions of interest for data collection. Tilt-series were collected at a nominal 42,000× magnification resulting in a calibrated pixel size of 3.37 Å or 3.45 Å, over a tilt range −40° to 64° for a positive pre-tilt with 3° increment, a total dose of ~140 e-/A^2^ and a nominal defocus range of −2 to −4.5 μm. Data acquisition was automated using a modified version of the dose-symmetric scheme (Hagen et al., 2017) taking the lamella pre-tilt into account and controlled using SerialEM (Mastronarde, 2005). For each tilt-series the CTF was estimated using gCTF (Zhang, 2016) and corrected for dose exposure (Grant and Grigorieff, 2015) using MATLAB scripts adapted for tomographic tilt-series (Wan et al., 2017). Tilt series alignment was performed using IMOD software package 4.9.2. (Kremer et al., 1996) by tracking fiducial markers spontaneously generated during the final sputtering in the Aquilos.

### Subtomogram averaging of NPC and diameter measurements

Subtomogram averaging was performed with slight modifications from a previously described workflow (Beck et al., 2007) using novaSTA package (https://github.com/turonova/novaSTA). Around 250 tomograms were reconstructed from 4× binned tilt-series. The reconstruction was done in IMOD using the SIRT-like filter to improve the contrast for manual picking of NPCs. In total 99 NPCs were picked. Subsequently the tomograms for subtomogram averaging were reconstructed with simple radial filter and 3D-CTF correction using novaCTF (Turoňová et al., 2017). The subtomogram averaging was performed using 8-fold symmetry on subtomograms which were down-sampled 8 times, followed the alignment on 4 times down-sampled subtomograms. Subsequently, the NPCs were split into asymmetric subunits with the subunits outside the lamellae being removed from the subsequent processing. The remaining set of 656 subtomograms was used to perform subtomogram averaging independently for the cytoplasmic, nuclear and inner ring. Fourier shell correlation at threshold 0.143 was computed for each ring and the respective resolutions were 36.3 Å, 35.8 Å, and 39.9 Å. Measurements of NPCs diameter from outer-inner nuclear membrane fusions and inner ring were carried out using inhouse MATLAB scripts.

### The distributions of hexamers in the capsid-like structures

Around 250 tomograms were reconstructed using 4× down-sampled tilt-series. The reconstruction was done in IMOD (Kremer et al., 1996). The SIRT-like filter option was used in order to generate tomograms with sufficient contrast for manual picking of viral structures.

Capsid-like structures identified in the cellular landscape were manually segmented using the IMOD drawing tool. The segmentation was used to create a surface in MATLAB which was represented as a set of triangles. The surface area of each capsid was then computed as a sum of areas of these triangles. For subtomogram averaging, a new set of 3D-CTF-corrected tomograms was reconstructed using novaCTF and downsampled 2×. Subtomogram averaging followed the protocol described in Mattei et al. (Mattei et al., 2016). The segmentation of virions was used to generate starting positions and the EMD-3465 map was resampled to match the data pixel size and used as an initial reference. Three iterations of alignment were run to allow the oversampled starting positions to shift to the positions corresponding to the hexamers in the CA lattice. Misaligned subtomograms were removed based on their angular deviation from the capsid shape and low cross-correlation values measured towards the reference. The cleaned set of subtomograms was used to generate a new reference. The whole subtomogram averaging procedure was run again from the oversampled starting positions using the new reference. After the removal of misaligned particles, the recovered lattice contained more CA hexamers. The visual representation of the lattice was created in UCSF Chimera (Pettersen et al., 2004) using Place Object plug-in (Qu et al., 2018).

